# Synaptic weight dynamics underlying memory consolidation: implications for learning rules, circuit organization, and circuit function

**DOI:** 10.1101/2024.03.20.586036

**Authors:** Brandon J Bhasin, Jennifer L Raymond, Mark S Goldman

## Abstract

Systems consolidation is a common feature of learning and memory systems, in which a long-term memory initially stored in one brain region becomes persistently stored in another region. We studied the dynamics of systems consolidation in simple circuit architectures with two sites of plasticity, one in an early-learning and one in a late-learning brain area. We show that the synaptic dynamics of the circuit during consolidation of an analog memory can be understood as a temporal integration process, by which transient changes in activity driven by plasticity in the early-learning area are accumulated into persistent synaptic changes at the late-learning site. This simple principle naturally leads to a speed-accuracy tradeoff in systems consolidation and provides insight into how the circuit mitigates the stability-plasticity dilemma of storing new memories while preserving core features of older ones. Furthermore, it imposes two constraints on the circuit. First, the plasticity rule at the late-learning site must stably support a continuum of possible outputs for a given input. We show that this is readily achieved by heterosynaptic but not standard Hebbian rules. Second, to turn off the consolidation process and prevent erroneous changes at the late-learning site, neural activity in the early-learning area must be reset to its baseline activity. We propose two biologically plausible implementations for this reset that suggest novel roles for core elements of the cerebellar circuit.

**Significance Statement:** How are memories transformed over time? We propose a simple organizing principle for how long term memories are moved from an initial to a final site of storage. We show that successful transfer occurs when the late site of memory storage is endowed with synaptic plasticity rules that stably accumulate changes in activity occurring at the early site of memory storage. We instantiate this principle in a simple computational model that is representative of brain circuits underlying a variety of behaviors. The model suggests how a neural circuit can store new memories while preserving core features of older ones, and suggests novel roles for core elements of the cerebellar circuit.

## Introduction

Memory systems transform transiently present information into a more persistent form. In short-term (working) memory, transient spiking activity of input neurons is transformed into persistent activity in downstream short-term memory-storing circuits (Zylberberg & Strowbridge 2017; Goldman-Rakic 1995). In long-term memory, memories stored transiently through neural plasticity at one site may become stored persistently at another site, through a process known as systems consolidation (Dudai *et al*. 2015; Squire *et al*. 2015). Systems consolidation is a common feature of learning and memory systems, including declarative memory (Genzel & Wixted 2017), fear conditioning (Do Monte *et al*. 2016), and motor skill learning (Krakauer & Shadmehr 2006). It is thought to help memory systems navigate the “stability-plasticity” dilemma (Abraham & Robins 2005; Grossberg 1987)—balancing the need to have capacity for new memories and the tendency of new memories to overwrite old ones, which could lead to catastrophic forgetting (McClelland *et al*. 1995; Roxin & Fusi 2013).

In working memory, the transformation of transient representations into a more persistent form can be characterized by well-established computational principles (Brody *et al*. 2003; Chaudhuri & Fiete 2016; Goldman *et al*. 2009; Major & Tank 2004; Seung 1996; Wang 2001; Zylberberg & Strowbridge 2017). For the storage of analog (graded or continuous-valued) memories, this transformation can be accomplished through temporal integration of transient activity in the input to a circuit into persistent changes in circuit output, with the set of possible stored activity patterns forming a continuous set of stable patterns.

For systems consolidation, overarching computational principles describing the transformation of transient into persistent representations are less well established. Qualitatively, the standard view of systems consolidation suggests that transient plasticity in an early-learning brain area results in altered neural activity that then triggers the induction of persistent changes in the late-learning area post-training (Do Monte *et al*. 2016; Krakauer & Shadmehr 2006; Hardt *et al*. 2013; Lesburguères *et al*. 2011; Richards & Frankland 2017; Squire *et al*. 2015). Through this process, the expression of learning becomes robust to inactivation of the early-learning area (Fig. 1*A,B*).

**Figure 1:**
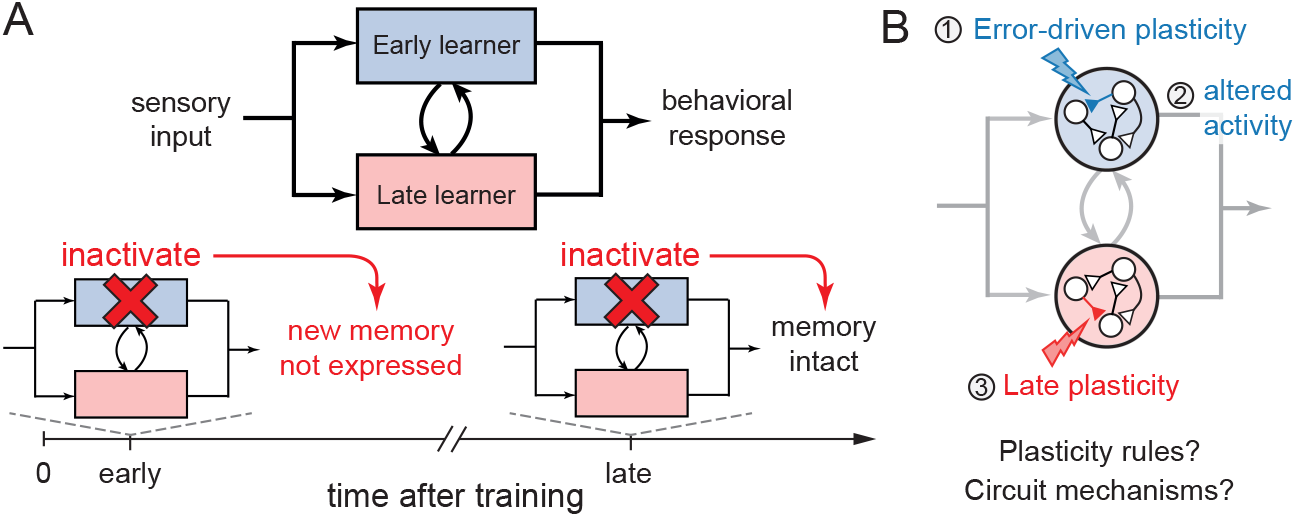
Schematic of systems consolidation. (*A*) Learned changes in the response to a sensory input occur first in an early-learning area, but are transferred over time to a late-learning area that becomes the sole site required for expression of learning, as revealed by inactivation experiments. (*B*) Feedback about behavioral errors drives synaptic plasticity in the early-learning area (1), leading to changes in activity (2) that in turn drive secondary plasticity in the late-learning area (3). Here, we investigate the dynamical mechanisms underlying this consolidation process and their implications for circuit organization and function.

We investigated the dynamics of systems consolidation in a model of a simple circuit that captures essential features of the systems consolidation of error-driven learning in brain areas such as the cerebellum (Cooke *et al*. 2004; Lisberger 2021; Raymond & Medina 2018), striatum (Andalman & Fee 2009; Warren *et al*. 2011; Yin *et al*. 2009; Makino *et al*. 2016; Teşileanu *et al*. 2017; Murray & Escola 2020), and amygdala (Medina *et al*. 2002; Do Monte *et al*. 2016). Building upon previous models of systems consolidation of oculomotor learning (An *et al*. 2023; Clopath *et al*. 2014; Herzfeld *et al*. 2020; Medina & Mauk 1999; Menzies *et al*. 2010; Porrill & Dean 2007; Yamazaki *et al*. 2015), we show that systems consolidation can be framed as a process of temporal integration, in which transient changes at the initial site of plasticity are integrated into changes at the final site. Further, within the context of cerebellar systems consolidation, our results extend previous proposals for how molecular layer interneurons and nucleo-olivary pathways, core circuit elements not included in traditional models, may serve to regulate cerebellar cortical plasticity (Herzfeld *et al*. 2020; Medina & Mauk 1999; Kenyon *et al*. 1998). Specifically, we show how these pathways may be necessary for stabilizing the dynamics of systems consolidation.

## Results

### A simple circuit model of systems consolidation

We modeled a simple circuit that learns the analog-valued gain of an input-to-output transformation. This gain is determined jointly by a direct and an indirect pathway through the circuit. Errors in behavioral performance drive plasticity at an early-learning site in the indirect pathway, which is then consolidated at a late-learning site in the direct pathway.

To make the analysis more concrete, we consider the specific example of oculomotor learning. Our focus is not on the intricacies of oculomotor learning specifically but the principles of consolidation that recur in circuit architectures throughout the brain. Nevertheless, our model captures many important features of oculomotor learning. The gain of eye movement responses to vestibular or visual stimuli can be adaptively modified by learning so as to attain any value within an analog range (Fig. 2*A*; see Materials and Methods, *Feedforward sensorimotor circuit model*; Broussard & Kassardjian 2004). Expression of this learned change in the sensory-to-motor transformation initially depends on the cerebellar cortex, but becomes cerebellum-independent within 24 hours post-training (Anzai *et al*. 2010; Jang *et al*. 2020; Kassardjian *et al*. 2005; Nagao & Kitazawa 2003; Shutoh *et al*. 2006). Learning and consolidation occur over timescales that are long compared to the eye movement responses to sensory stimuli, hence we modeled the latter as instantaneous.

**Figure 2:**
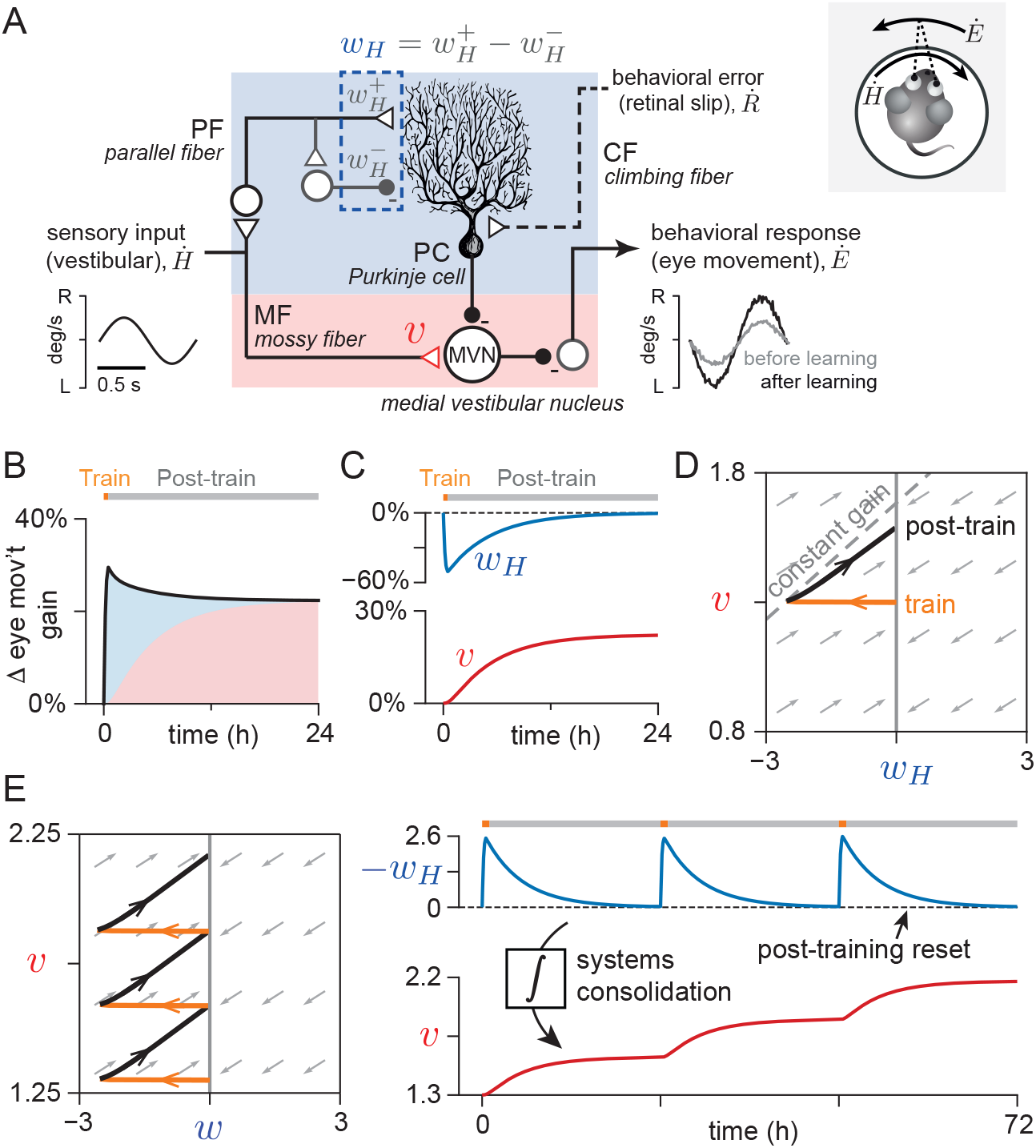
Synaptic weight dynamics in a representative feedforward-architecture circuit model of systems consolidation correspond to a temporal integration process. (*A*) The microcircuit underlying oculomotor learning alters the amplitude (“gain”) of the reflexive eye movement response to sensory (vestibular) input. Incorrect eye movement amplitude causes image motion on the retina (“retinal slip”). Instructive signals carrying information about behavioral errors, i.e., retinal slip, drive changes in weight, *w*_*H*_, at an early-learning site in the cerebellar cortex (blue shaded area), with *w*_*H*_ representing the difference between excitatory 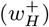 and inhibitory 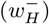 synaptic strengths. Over time, expression of learning becomes dependent only on the weight *v* at a late-learning site in the brainstem (red shaded area). Excitatory synapses are represented by open triangles, inhibitory synapses by filled circles. (*B, C*) Simulation of model for 30 minutes of training (orange block) to increase the input-to-output gain of the eye movement response, followed by 23.5 hours post-training in the dark (grey block). During the post-training period, the model received no information about errors. (*B*) Change in the gain of the eye movement response (black line), with shading showing contributions from the early-(blue) and late-learning (red) sites. (*C*) Change in weights *w*_*H*_ (blue) and *v* (red). (*D*) Trajectory of synaptic weights during the training (orange) and post-training (black) periods. Grey arrows show the analytically calculated, approximate instantaneous direction in which a weight configuration at a given point will evolve without information about errors. All trajectories tend toward some marginally stable point along the line *w*_*H*_ = 0 (thick grey line). A trajectory resulting in perfect consolidation would follow the “constant gain” line (grey dashed line). (*E*) Consolidation corresponds to a temporal integration process in which transient changes in activity at the early-learning site, driven by plasticity at *w*_*H*_, are accumulated into persistent changes in *v*. Panels show trajectory and dynamics in synaptic weight space (*left*) and over time (*right*) in a simulation of three consecutive days of the training protocol in *B–D*.

We first studied a model of the circuit in which the input-to-output transformation was purely feedforward. In the context of cerebellum-dependent learning, this is consistent with the classical Marr-Albus-Ito model (Albus 1971; Marr 1969; Ito 1982) and previous computational studies of the consolidation of oculomotor learning (Clopath *et al*. 2014; Herzfeld *et al*. 2020; Porrill & Dean 2007; Yamazaki *et al*. 2015). We simulated neural activity and changes in synaptic weights in the feedforward circuit model during a training and post-training period. During the training period, the model receives instructive signals about behavioral performance, which control the induction of plasticity at the early-learning site. For the case of oculomotor learning, these instructive signals reflect behavioral errors resulting from failures of eye movements to stabilize images on the retina, carried by the climbing fiber input to the cerebellar cortex. Such signals induce a decrease in the weight *w*_*H*_ at the early-learning site, reducing the inhibitory output from the indirect (cerebellar cortical) pathway and thereby increasing the overall gain of the sensory-to-motor transformation (Fig. 2*B,C* ; Boyden *et al*. 2004; Inoshita & Hirano 2018; Jang *et al*. 2020; Kimpo *et al*. 2014).

During the subsequent post-training period, information about behavioral errors is not available (experimental subject is placed in the dark to eliminate visual feedback about the stabilization of images on the retina), and we assume *w*_*H*_ decays back to a baseline value of zero (Fig. 2*C*). This baseline value of zero for *w*_*H*_ can be interpreted as a balance of excitation and inhibition at the early-learning site. The decay of plasticity at *w*_*H*_ is consistent with electrophysiological measurements (Jang *et al*. 2020), the experimental observation that memory expression becomes cerebellum-independent over time (Anzai *et al*. 2010; Kassardjian *et al*. 2005; McElligott *et al*. 1998; Nagao & Kitazawa 2003), and previous models (An *et al*. 2023; Clopath *et al*. 2014; Herzfeld *et al*. 2020; Yamazaki *et al*. 2015).

Because the synaptic changes induced at the early-learning site during the training period are transient, successful consolidation requires the induction of persistent changes at the late-learning site (An *et al*. 2023; Clopath *et al*. 2014; Herzfeld *et al*. 2020; Medina & Mauk 1999; Menzies *et al*. 2010; Porrill & Dean 2007). Consolidation is known to depend on neural activity during the post-training period (Okamoto *et al*. 2011), which presumably induces the plasticity at the late-learning site (Jang *et al*. 2020). To achieve this, we implemented a heterosynaptic plasticity rule for the late-learning weight *v* of the form

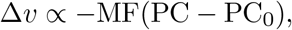

in which weight changes are driven by the correlation of direct pathway input (in the cerebellar context: mossy fiber, MF) with early-learning area output (Purkinje c ell, PC) relative to baseline (PC_0_) (see Materials and Methods, *Learning rules*, for the full equation), consistent with previous modeling of oculomotor learning (An *et al*. 2023; Clopath *et al*. 2014; Herzfeld *et al*. 2020; Medina & Mauk 1999; Menzies *et al*. 2010; Porrill & Dean 2007). As a result, altered post-training activity at the early-learning site induced an increase in *v* to a new steady-state value (Fig. 2*C*). This change persisted even as *w*_*H*_ returned to its baseline, supporting a persistent increase in the input-to-output gain of the circuit (Fig. 2*B,C*).

### Dynamical principles of systems consolidation

To understand the features of the synaptic weight dynamics that support successful systems consolidation, we plotted the trajectory describing the joint evolution of the early- and late-learning weights. Initially, training induces a change at the early-learning site, decreasing the weight *w*_*H*_ from its baseline value and moving the weights (*w*_*H*_, *v*) (Fig. 2*D*, orange trajectory) to a point corresponding to an increased input-to-output gain of the eye movement response (Fig. 2*D*, dashed “constant gain” line). During the post-training period, the early-learning weight decays to baseline, but the late-learning weight is driven to a new steady state value that preserves some of the increase in the input-to-output gain accrued during training (Fig. 2*D*, black trajectory). A nearly parallel trajectory will be followed by the evolution of the weights during the post-training period for any weight configuration reached during training (Fig. 2*D*, grey arrows), with all trajectories approaching some steady state value along a line in synaptic weight space (Fig. 2*D*, dark vertical line). Because the weight dynamics are at steady state whenever the activity of the early-learning area is at baseline, which occurs when the early-learning weight *w*_*H*_ decays to zero, there is a continuum of values that can be stably maintained by the late-learning weight *v* (see SI Text, S1). Therefore, for a given initial value of *v* before training, the final value reached after consolidation varies in a graded manner with the magnitude of the change in the early-learning weight *w*_*H*_ during training. This enables the circuit to learn and maintain any graded amplitude input-to-output gain across multiple learning events (Fig. 2*E*, S1; see SI Text, S1).

These dynamics suggest an intuitive computational principle: systems consolidation of analog memories corresponds to a temporal integration process in which the late-learning weight stably accumulates changes induced during training at the early-learning site. For the simple feedforward model we have been examining, this can be visualized directly in the space of synaptic weights as an integration of transient changes in *w*_*H*_ into persistent changes in *v* (Fig. 2*E*). More generally, changes in the output of the early-learning area, driven by plasticity at *w*_*H*_, are accumulated into weight changes at the late-learning site (see below for more complex circuit architectures).

To obey this principle, two conditions must be satisfied. First, the rule governing plasticity at the late-learning site must support the stable accumulation of persistent weight changes and corresponding continuum of input-to-output gains. Second, the circuit must reset the output of the early-learning site post-training so that the accumulation in the late-learning weight stops. Here we lay out the implications of the principle of consolidation as integration, starting with circuit function. Then, we show how the two simple conditions stated above place strong constraints on the plasticity rules and circuit features needed to support systems consolidation.

### Consolidation exhibits diffusive drift in the absence of information about errors, suggesting a speed-accuracy tradeoff for consolidation

Because the dynamics of systems consolidation can be understood as temporal integration, systems consolidation exhibits properties that have been well-characterized in other kinds of integrators, such as diffusive drift in the value being memorized due to the accumulation of noise, which has been found in neural integrator circuits implementing working memory (Brody *et al*. 2003; Burak & Fiete 2012; Ganguli *et al*. 2008; Lim & Goldman 2012; Seung 1996). In our model, noise in the neural activity driving plasticity at the late-learning weight *v* is accumulated, leading the input-to-output gain of the circuit to drift. In the absence of information about behavioral errors (e.g. during the post-training period), even if the output of the early-learning area is reset on average, noise in the early-learning weight *w*_*H*_—for example, resulting from noise in the pathway that carries instructive signals during learning—is integrated into changes in *v* (Fig. 3*A–D* ; see SI Text, S1, for analytical derivation of how the noise accumulates). The amplitude of changes in *v* post-training, resulting from changes in *w*_*H*_, is defined by the slope of the flow field (Fig. 3*A,B*, grey arrows), which is proportional to the learning rate of *v* and inversely proportional to the rate at which the early-learning site *w*_*H*_ is reset to baseline. This implies that a system with fast consolidation will reach a given level of consolidated learning quickly (i.e., in a small number of training sessions) but accumulate noise quickly as well. By contrast, a system with slow learning will require many training sessions to reach the same level of consolidated learning, but will reach this level with a smaller total level of consolidated noise (Fig. 3*E–H*). This represents a type of “speed-accuracy” tradeoff in achieving a given level of consolidated learning, similar to that seen in integration-based tasks like accumulation of evidence for decision-making (Bogacz *et al*. 2010; Gold & Shadlen 2007).

**Figure 3:**
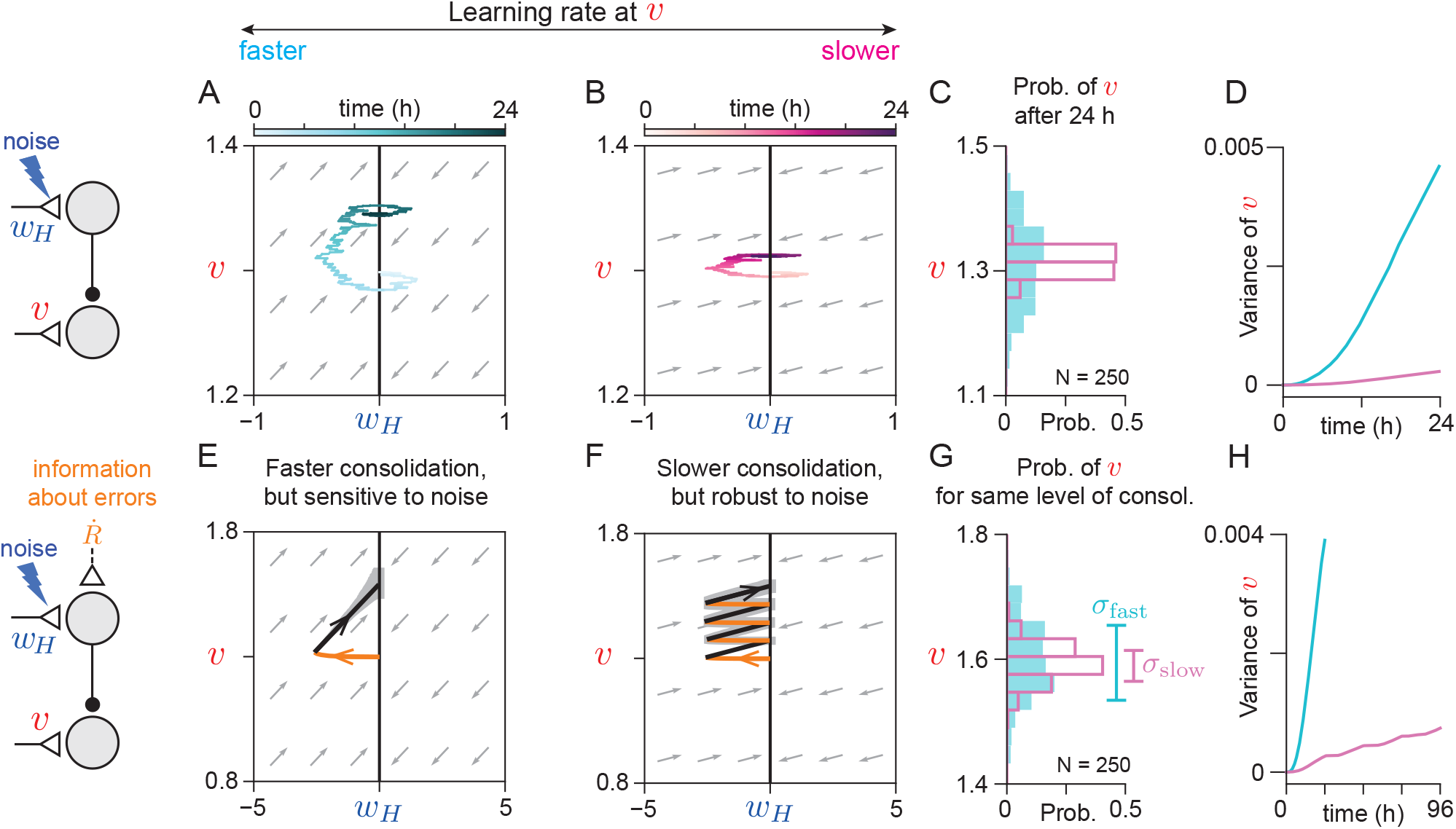
Diffusive drift of consolidated memory implies a tradeoff between speed of consolidation and sensitivity to noise. (*A, B*) Trajectories of synaptic weights in response to periodic random perturbations in the early-learning weight *w*_*H*_ when the learning rate at *v* is relatively fast (*A*) or slow (*B*) (light to dark colors show progression of time). (*C*) Distribution of the value of the late-learning weight *v* at the end of 24 hours of random perturbations of *w*_*H*_ for *N* = 250 simulations (see Materials and Methods for details), using the slower (open magenta bars) or faster (filled cyan bars) learning rates at *v* depicted in panels *A* and *B*, respectively. (*D*) Time course of the variance of the distributions in *C*. (*E, F*) Mean trajectories of synaptic weights during training (orange) and subsequent post-training periods (black) with same perturbations as in *A* and *B* (*N* = 250 simulations). Grey band indicates the range of trajectories within one standard deviation of the mean at each time point. The learning rates in panels *E* and *F* were the same as in panels *A* and *B*, respectively. Trajectories in panel *F* reach approximately the same mean level of consolidation as those in panel *E* after four consecutive days of training. (*G*) Distribution of *v* at the end of training for simulations shown in *E* (filled cyan bars) and *F* (open magenta bars). Vertical bars show standard deviations of the distributions, *σ*_fast_ and *σ*_slow_ respectively. (*H*) Time course of the variance of *v* due to perturbations for simulations shown in *E* (cyan) and *F* (magenta). With a slower learning rate at *v*, the circuit accumulates less noise. The scalloping in the variance for the simulations in *F* is due to the effect of feedback about errors during each training session.

### Consolidation mitigates the plasticity-plasticity dilemma through averaging

Framing consolidation as integration highlights mechanistically how systems consolidation can address the stability-plasticity dilemma (Abraham & Robins 2005; Grossberg 1987), i.e., the tradeoff between the ability of neural systems to adapt quickly and to not overwrite previous learning. Within the context of error-driven learning, this tradeoff arises when the optimal, “target” input-to-output gain of the circuit transformation, as conveyed by the instructive signals guiding learning, fluctuates across training sessions due to changes in the environment or in the motor plant (e.g., due to experimental manipulations, fatigue, injury, or changes in body mass). With one site of learning, a more plastic circuit that adapts quickly to reduce error during a given training session will tend to have a bigger initial error at the start of the next training session (Fig. 4*A–C*, “*w*_*H*_ fast”). On the other hand, a more stable circuit, whose gain adapts slowly and approximates the mean of the fluctuating learning target, will have, on average, smaller initial errors across training sessions, but will less fully reduce the error during a given training session (Fig. 4*A–C*, “*w*_*H*_ slow”; SI Text, S2).

**Figure 4:**
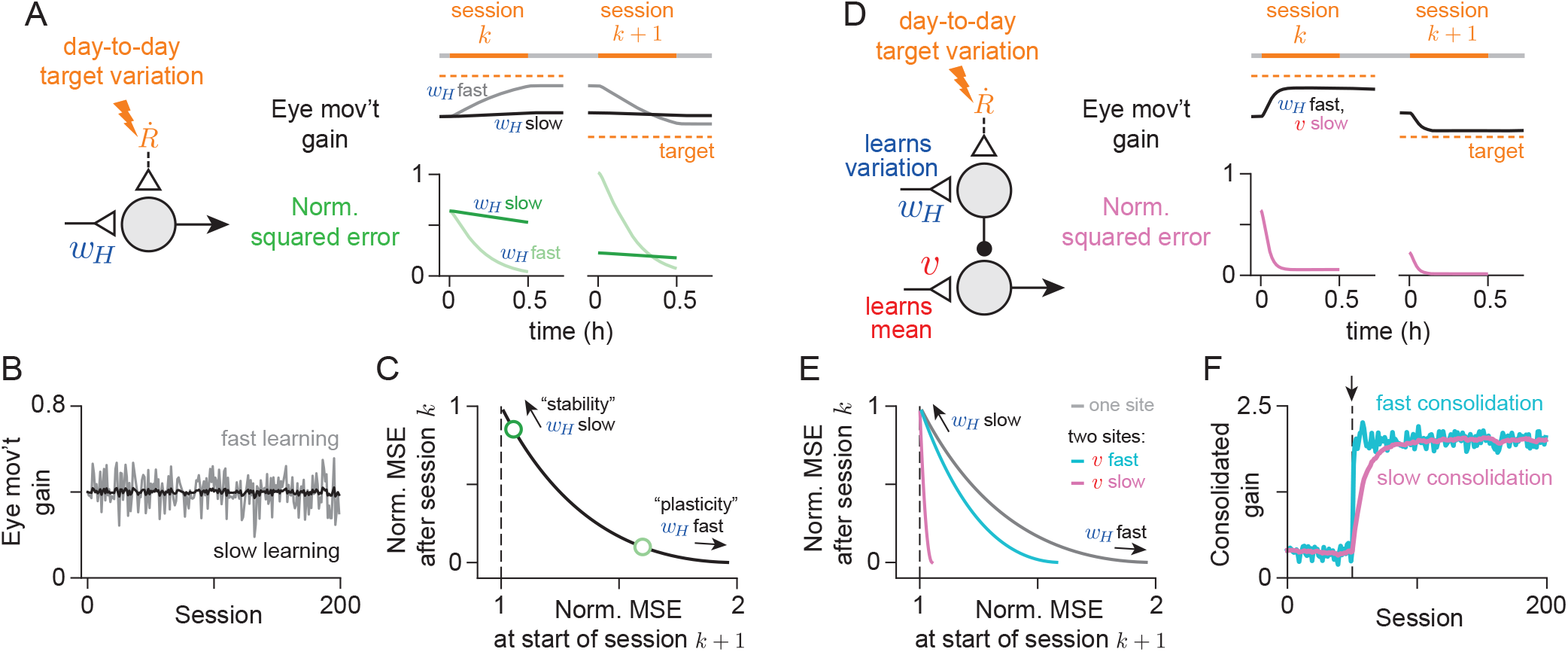
Consolidation averages over variability in the instructive signal to mitigate the stability-plasticity dilemma. (*A*) Learning in a model of a circuit with one site of plasticity that performs a graded amplitude input-to-output transformation is subject to the stability-plasticity dilemma. During a training session, the instructive signal drives a change in the weight *w*_*H*_, which is stably remembered post-training. Across training sessions, the externally instructed target value of the input-to-output gain varies randomly about a mean value. Fast learning at *w*_*H*_ (light colors) reduces error more effectively during a training session compared to slow (dark colors), but increases the size of the expected error at the beginning of the next training session (distance from orange dashed line). (*B*) Input-to-output gain over 200 training sessions in a simplified model that simulates weight changes discretely over sessions (SI Text, S2). Slower learning (black) results in a more stable value of the weight due to greater averaging over variability in the instructive signal. (*C*) Expected (mean) squared error (MSE) after a training session in the simplified model trades off with expected error at start of subsequent session, calculated analytically and normalized to the variance of the target gain distribution. Dark and light green circles correspond to the learning rates used to generate black and grey curves in panel *B*. (*D*) A circuit with two sites of learning can mitigate the tradeoff in panel *C*. A persistent, slow-learning site of consolidation, *v*, can estimate the mean of the expected gain distribution, while a forgetful, fast-learning early site, *w*_*H*_, can account for day-to-day variation in the instructive signal. (*E*) Relationship between normalized expected MSE at the end versus start of a training session when there are two sites of learning with fast (cyan) or slow (magenta) learning at the late site *v*.Grey curve replots the relationship when there is only one site of learning (same as *C*). (*F*) Consolidated gain in response to a change in the mean of the target gain distribution (at session indicated with arrow) for relatively fast (cyan) or slow (magenta) learning at *v* (rates same as *D*, but using the simplified model). Slow consolidation leads to reduced variability in the long run, but at the cost of larger errors early in training.

With two sites of learning—a fast-learning and fast-forgetting early site of plasticity, and a slow-learning late site of consolidation—the circuit can mitigate this tradeoff. Slow consolidation at *v* stores the long-term average of the fluctuating target gain conveyed by the instructive signals across training sessions, while fast plasticity at *w*_*H*_ adapts quickly during a single training session but is reset post-training (Fig. 4*D*). In this way, the circuit can both minimize the average error at the start of each training session and respond quickly to reduce errors within a session (Fig. 4*E* ; SI Text, S2). However, even with two sites of learning, the minimization is not perfect, and there is a tradeoff in the size of the expected error at the start of the next training session versus the number of sessions required to consolidate the mean target gain (Fig. 4*F*). This is another form of the speed-accuracy tradeoff discussed in the previous section (Fig. 3).

### Implications for plasticity rules

To support consolidation, the late-learning site must be able to stably accumulate persistent weight changes over time. This requires that the plasticity rule at the late-learning site support a continuum of stable weight values, so that the synapse can persistently hold any change in the weight. This is readily achieved by a heterosynaptic plasticity rule (Fig. 2*E*), because weight changes stop whenever the activity at the early-learning site returns to its baseline or, more generally, becomes uncorrelated with the direct pathway input (Dean *et al*. 2002).

Recent work has alternatively proposed a Hebbian, covariance-like rule for the consolidation of oculomotor learning (Yamazaki *et al*. 2015). Here we considered a Hebbian covariance-like rule of the form

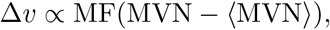

in which plasticity is proportional to presynaptic activity (mossy fiber, M F) multiplied by the difference between postsynaptic activity (medial vestibular nucleus, MVN) and a sliding threshold equal to its recent average (angle brackets) (see Materials and Methods, *Learning rules*, for the full equations). Our analysis shows that this covariance-like rule can support a continuum of stable values, and hence consolidation, if the circuit receives no time-varying input or noise during the post-training period (Fig. 5*A,B*, light lines). However, in the more biologically realistic case that the circuit does receive time-varying input or noise, the synaptic weight *v*, and thus the input-to-output gain, becomes unstable, growing exponentially during the post-training period (Fig. 5*A,B*, dark lines; SI Text, S3).

**Figure 5:**
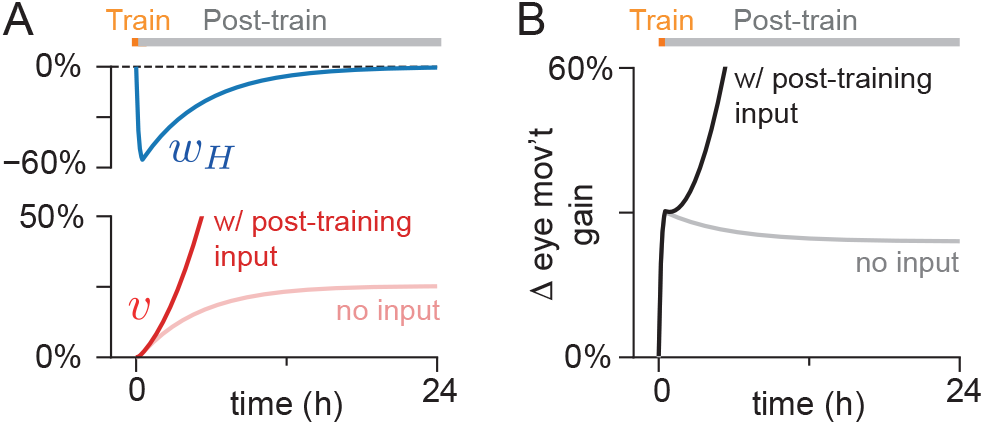
Hebbian covariance rule cannot support stable consolidation in the presence of time-varying signal or noise. (*A*) Evolution of the early- and late-learning weights, *w*_*H*_ (blue) and *v* (red), over time when using a Hebbian covariance rule at *v*, during a simulation of 0.5 h of training to increase the input-to-output gain of the circuit, followed by 23.5 h post-training without information about errors, when time-varying input to the circuit was either present (dark lines) or not present (light “no input” lines) post-training. (*B*) Change in the input-to-output gain corresponding to the simulations in *A*.

The failure of the covariance-like rule reflects an inherent source of instability in Hebbian learning (Miller & MacKay 1994). In the basic Hebbian rule, correlations between presynaptic and postsynaptic firing rates drive increases in synaptic weights. These increased synaptic weights in turn drive postsynaptic activity and thus increased correlations between pre- and postsynaptic firing, forming a positive feedback loop. For the covariance-like rule considered above, the sliding threshold counters the increased correlations associated with changes in the *average* postsynaptic firing rate, but does not counter the increased correlations associated with *fluctuations around* the average (see SI Text, S3; Loewenstein 2008). Other proposed methods for countering this instability, including weight normalization (Miller & MacKay 1994; Oja 1982) and firing rate-target homeostasis (Bienenstock *et al*. 1982; Chistiakova *et al*. 2015; Turrigiano 2008; Yger & Gilson 2015), also fail to support a continuum of stable values in our model. Rather, they yield dynamics that drive the late-learning weight to a single stable value (see SI Text, S4). Altogether, this analysis suggests that either heterosynaptic plasticity, or a different variant of Hebbian rule from those typically considered, is required to implement systems consolidation of an analog memory.

### A circuit with an internal feedback loop requires an active post-training reset mechanism at the early-learning site

In addition to constraints on the form of plasticity at the late-learning site, consolidation also requires that the output of the early-learning area be reset during the post-training period, stopping the integration at the late-learning site. For the simple feedforward model described above, this condition is met by the intrinsic decay of the early-learning weight *w*_*H*_ to zero, which represents a balance of excitatory and inhibitory input to the early-learning area.

An active, rather than passive, post-training reset mechanism is necessary for stable consolidation in the more complex case where the input-to-output transformation computed by the circuit contains an internal feedback loop (Fig. 6*A*). We modeled this case by adding to the model an internal feedback connection from the late-to the early-learning area. For oculomotor learning, the internal loop corresponds to an efference copy of the eye movement command (Miles & Lisberger 1981; Lisberger & Sejnowski 1992; Lisberger 1994; Person 2019). Here we describe a model with a fixed internal feedback weight *w*_*E*_ *>* 0; similar results are obtained if *w*_*E*_ is plastic (Fig. S2; SI Text, S6). As in the feedforward model, the condition that plasticity at the late-learning site support stable accumulation can be met by a heterosynaptic plasticity rule at *v*. However, the reset condition can no longer be met just by the passive decay of the weight *w*_*H*_ of the feedforward pathway in the early-learning area to a fixed baseline, but rather requires that, during the post-training period, *w*_*H*_ returns to a value that depends on *v*. This is because, due to the internal feedback from the late-to the early-learning area, changes in *v* also drive altered activity of the early-learning area. These changes must be offset post-training by changes in *w*_*H*_ so the feedforward sensory input and the feedback from the late-learning area effectively cancel, eliminating the drive for plasticity at the late-learning site and enabling *v* to reach a new steady state.

**Figure 6:**
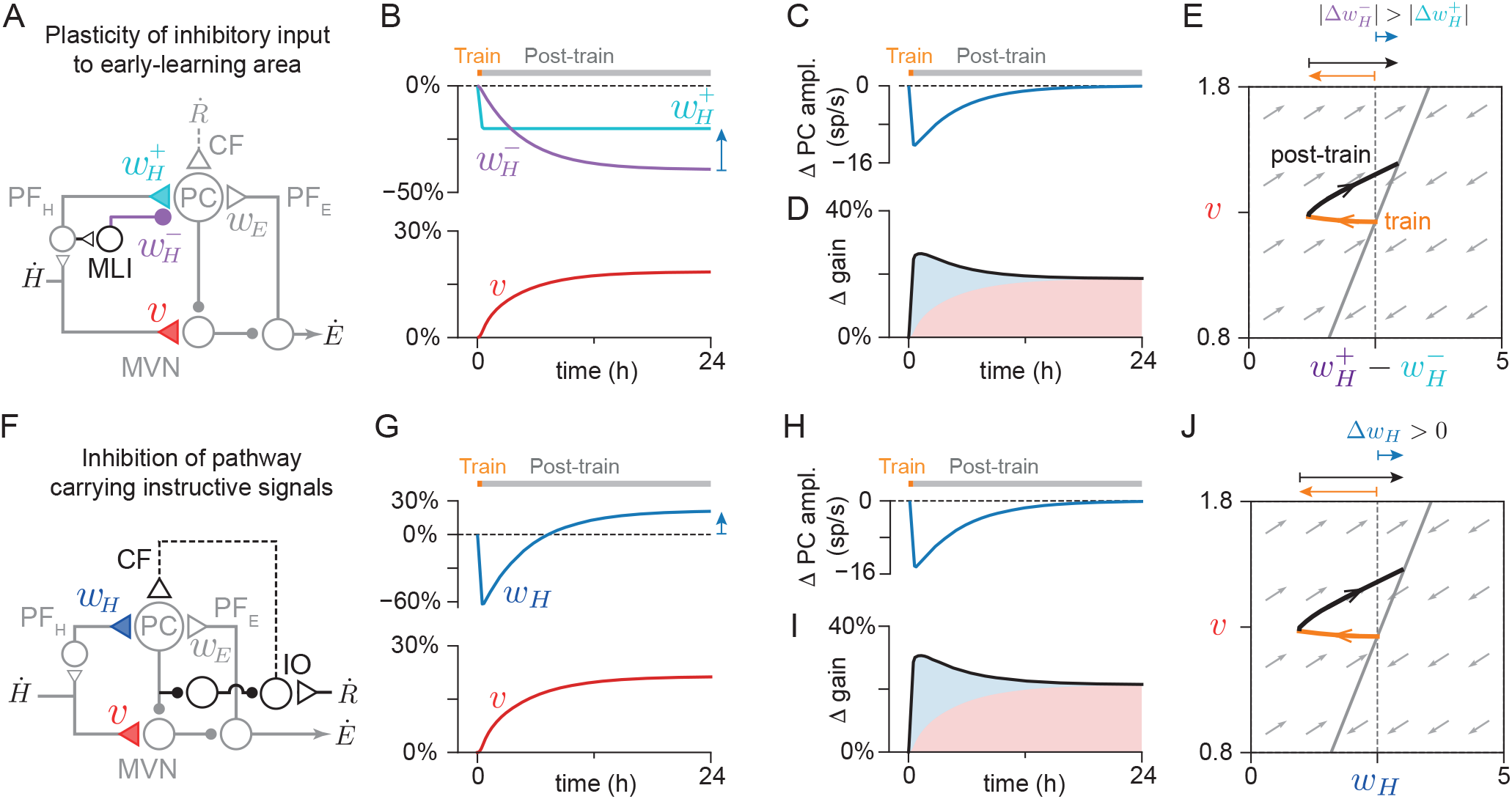
Circuit mechanisms for post-training reset enable consolidation in the presence of an internal feedback loop. (*A*–*E*) Reset of early-learning area output via plasticity of inhibitory input onto the early-learning area. (*A*) Circuit diagram. PF_*H*_ and PF_*E*_ : cerebellar parallel fiber inputs to Purkinje cells (PC) carrying sensory (vestibular) input 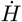 and efference copy feedback of motor output *Ė*. MLI: molecular layer interneuron. CF: climbing fiber. 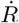 : instructive signal carrying information about behavioral (retinal slip) errors (*B*) Change in early-learning excitatory weight 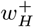 (cyan), inhibitory weight 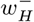 (purple) and late-learning weight *v* (red), during 0.5 h of simulated training (orange block) to increase the input-to-output gain of the circuit, followed by 23.5 h post-training with no information about errors (grey block). At steady state, the value of 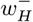 has decreased more than 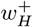, so that the net change in the net weight 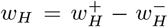 is positive (blue arrow). (*C*) Early-learning area output (amplitude of Purkinje cell activity relative to moving baseline), which drives consolidation at the late-site. (*D*) Change in the gain of the eye movement response (black line) to a sensory input. Blue and red shaded areas show the contribution of the early- and late-learning areas to the circuit transformation. (*E*) Trajectory of synaptic weights during the training (orange) and post-training (black) periods. During training, the net early-learning weight decreases (orange arrow above plot) from its initial value (dashed black line), but post-training (black arrow) approaches a steady state value that is larger than before training (blue arrow). Grey arrows show the approximate instantaneous direction in which a weight configuration at a given point in synaptic weight space will evolve during the post-training period, determined analytically. All trajectories tend toward a marginally stable point along a line (solid grey line). (*F* –*J*) Reset of early-learning area output via inhibition of the pathway carrying instructive signals to the early-learning site. (*F*) Circuit diagram. IO: inferior olive. (*G*) Change in early-learning weight *w*_*H*_ (dark blue) and late-learning weight *v* (red). The post-training reset drives *w*_*H*_ to a steady state value larger than before training (blue arrow). (*H–J*) Same as *C–E*, but for model shown in *F*.

### The reset requirement suggests a role for specific features of the cerebellar circuit architecture in consolidation

Active reset of the early-learning site could be achieved in at least two ways. First, the feedforward input to the early-learning area can be decomposed into the sum of a direct excitatory and disynaptic inhibitory pathway that have weights 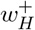 and 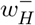, respectively, controlled by separate plasticity mechanisms (Fig. 6*A*). We assume that changes in the excitatory weight are driven only by instructive signals occurring during the training period, with no passive decay. We then find that, during the post-training period, plasticity in the inhibitory weight can reset the activity of the early learning area such that it no longer responds to sensory input. This resetting is achieved when the inhibitory input weight is governed by a Hebbian rule driven by correlations between presynaptic inhibitory inputs and postsynaptic spiking relative to a sliding threshold (Fig. 6*B–E* ; Materials and Methods, *Circuit model with internal feedback loop*; Vogels *et al*. 2011). Alternatively, the reset could be driven by inhibition from the late-learning area onto the source of the instructive signals that control plasticity at the early-learning site (Fig. 6*F–J* ; Materials and Methods, *Circuit model with internal feedback loop*). In the cerebellar context, the first reset mechanism would correspond to plasticity of molecular layer interneuron-to-Purkinje cell synapses, and the second could be achieved by the inhibitory pathway from the cerebellar nuclei to the inferior olive (see Discussion). Thus, a central computational role for these components of the cerebellar circuitry may be to implement the post-training reset of the rapid, early plasticity in the cerebellar cortex to support stable consolidation of learning.

Our model predicts that, following consolidation of learning to increase the input-to-output gain of the circuit, the sensitivity of cells in the early-learning area to feedforward input, *w*_*H*_, will be higher than the pre-training baseline, as has been suggested based on experimental measurements (Miles *et al*. 1980; Lisberger 1994). This can be understood more clearly by plotting the dynamics of the synaptic weights. During consolidation, the post-training reset dynamics cause the weights to move along a trajectory to a new steady state along a line in the space of synaptic weights. In the feedforward model, this line is vertical, so the early-learning weight returns to its pre-training baseline during the post-training consolidation period (Fig. 2*E*). When there is an internal feedback loop, the line has a finite, positive slope, so the net weight *w*_*H*_ does not just return to its pre-training baseline value, but goes to a steady state value that is larger than before training (Fig. 6*E,J*, black arrows). Thus, our model predicts that, after consolidation, the change in the early-learning weight, relative to pre-training, will be in the opposite direction from the change in weight during training (Fig. 6*E,J*, blue vs orange arrows above plot; Payne *et al*. 2024).

## Discussion

We propose a computational principle governing systems consolidation of analog memories: systems consolidation is defined by a temporal integration process in which the late-learning weight stably accumulates changes induced during training at the early-learning site. To obey this principle, the circuit must have certain properties: First, the synaptic weight at the late-learning site must be governed by plasticity rules that enable it to stably accumulate and maintain any one of a continuum of values. We show that this is achieved by a heterosynaptic plasticity rule but not by standard stabilized Hebbian rules. Second, the output of the early-learning area must be reset post-training to stop the accumulation of weight changes at the late-learning site.

The accumulation of synaptic weight changes underlying systems consolidation is reminiscent of the temporal integration of transient spiking activity into persistent spiking activity in neural circuits that accumulate and store information in working memory, such as that implicated in the accumulation and storage of neural signals encoding evidence in decision-making tasks (Brody *et al*. 2003; Chaudhuri & Fiete 2016; Churchland & Ditterich 2012; Gold & Shadlen 2007; Goldman *et al*. 2009; Major & Tank 2004; Usher & McClelland 2001; Wang 2001; Wang 2008; Zylberberg & Strowbridge 2017). Our model shares distinctive features with such neural integrator circuits, arising from their similar underlying dynamics. First, it is well known that integrator circuits require a fine tuning of parameters for stability of the memory (Chaudhuri & Fiete 2016; Goldman *et al*. 2009; Seung 1996; Zylberberg & Strowbridge 2017). This fine tuning occurs in two places: (1) in the strength of the recurrent connections between neurons, which offsets intrinsic decay of neural activity; and (2) in the strength of tonic background inputs entering the network. In the case of systems consolidation, we assume perfect integration (no decay) at the late-learning site, so only the second type of tuning is required. Specifically, output from the early-learning area must be reset post-training; otherwise, it will drive incessant weight changes at the late-learning site. This is analogous to the finding that neural integrator circuits will integrate tonic background spiking activity unless these inputs are subtracted out or otherwise nullified (Cannon *et al*. 1983; Goldman *et al*. 2009; Seung 1996). Drawing on the neural integrator literature, one proposed solution to this fine-tuning problem is to turn the continuum of steady states into a set of (many) finely discretized, robustly stable fixed points (Goldman *et al*. 2003; Koulakov *et al*. 2002). This could correspond to our single “lumped” synaptic weight *v* being composed of the sum of the binary weights of individual bistable synapses (O’Connor *et al*. 2005) that are recruited with different thresholds (Goldman *et al*. 2003; Nikitchenko & Koulakov 2008). Second, noisy input causes diffusive drift that progressively corrupts the stored memory (Fig. 3; Burak & Fiete 2012; Ganguli *et al*. 2008;

Lim & Goldman 2012). In our model, this led to a tradeoff between the speed of consolidation and the accuracy of the consolidated memory (Fig. 3; SI Text, S1). This can be interpreted as the long-term memory analog of the speed-accuracy tradeoff in decision-making tasks (Bogacz *et al*. 2010; Gold & Shadlen 2007).

The similarities in dynamics between neural integrator circuits and our systems consolidation model arise from the fact that in both cases, the memory stored in the circuit is analog—in our case, the amplitude of the circuit’s learned response to a given input is graded. Much of the previous theoretical work on systems consolidation has instead considered the case of effectively binary neurons (Alvarez & Squire 1994; Murray & Escola 2020; Remme *et al*. 2021; Roxin & Fusi 2013; Tomé *et al*. 2022; Wittenberg *et al*. 2002), in which memory expression is evaluated with respect to whether or not a given neuron fires, rather than the graded value of its firing rate. In such models, Hebbian plasticity typically drives synaptic weights to take either a high (saturated) or a low value, making individual synaptic dynamics effectively bistable (Dong & Hopfield 1992; Miller & MacKay 1994). By contrast, in our model, the synaptic weight at the late-learning site must be able to stably take any analog value within a continuum, which is naturally achieved by a heterosynaptic plasticity rule. Although such heterosynaptic rules are commonly used in modeling supervised learning tasks (Dayan & Abbott 2005), they may form a more specialized class of biological plasticity than Hebbian learning rules. Supporting the continuum of steady states required for consolidation is more challenging for classical Hebbian rules (see SI Text, S4), though may be possible with more complex forms of Hebbian plasticity, such as a three-factor (Kuśmierz *et al*. 2017) or dendritic plasticity rule (Urbanczik & Senn 2014). Alternatively, as noted in the previous paragraph, it is possible that the apparent continuum of weight values modeled here instead reflects a sum of discrete, bistable synaptic contributions.

The framing of systems consolidation as a process of temporal integration provides mechanistic insight into previous work on how consolidation addresses the stability-plasticity dilemma (Abraham & Robins 2005; Grossberg 1987). In our model, the stability-plasticity dilemma is mitigated because, in the presence of variability in the instructive signals about behavioral errors, slow integration (i.e., averaging) at the late-learning site tracks long-timescale behavioral requirements, while the early-learning site quickly tracks short-timescale changes in these requirements (Fig. 4*D* ; for a related use of integration to improve deep network training, see Johnson & Zhang 2013). In this manner, averaging leads to a more general memory at the late-learning site that is constantly updated by specific but transient memories at the early-learning site (Sekeres *et al*. 2018; Tse *et al*. 2007; Lindsey & Litwin-Kumar 2023; Richards & Frankland 2017; Sun *et al*. 2023). This framing could also be extended to describe synaptic consolidation, through which transient early plasticity is consolidated into persistent, graded late plasticity in the same synapse (Jacquerie *et al*. 2023; Leimer *et al*. 2019; Li & Van Rossum 2020). The timescale over which the late-learning site averages is defined by the relative rates of plasticity at the late- and early-learning sites (Fig. 4*F*). This may suggest that the rate at which the circuit consolidates (i.e., the fraction of learning consolidated post-training) is tuned so that the timescale of the average is matched to the timescale over which the mean is expected to change in the world (Körding *et al*. 2007).

Though systems consolidation appears to be a common feature of learning and memory systems, the details of how it occurs in each system may be shaped by specific computational needs. In hippocampus-dependent memory, the early-learning circuit of the hippocampus learns the associations between components or features of an episode. Post-training replay of the activity patterns representing these associations then drives consolidation to the neocortex (Carr *et al*. 2011; Squire *et al*. 2015; Alvarez & Squire 1994; Tomé *et al*. 2022; Roxin & Fusi 2013; Murre 1996; Wittenberg *et al*. 2002). In the system we model, the circuit learns associations between a sensory input and a behavioral error signal that enforces a desired output. Post-training “replay” of the activity patterns learned at the early site then drives consolidation of the learned input-to-output transformation at the late-learning site. Although in both cases the correlations present during training are recapitulated in the post-training neural activity, hippocampal replay occurs as discrete events with transient, spontaneous reactivation of representations (e.g., during a sharp wave ripple), whereas in the cerebellum the “replay” could be driven by ongoing input to the circuit.

Our work extends previous modeling of the consolidation of oculomotor learning (An *et al*. 2023; Clopath *et al*. 2014; Herzfeld *et al*. 2020; Medina & Mauk 1999; Porrill & Dean 2007; Yamazaki *et al*. 2015) in two key ways. First, we analyzed the conditions under which consolidation can occur successfully. We showed that consolidation requires that plasticity at the late-learning site support a continuum of steady state weight values, a requirement readily met by a heterosynaptic plasticity rule. This explains why heterosynaptic rules have been successful in previous work (Clopath *et al*. 2014; Herzfeld *et al*. 2020; Medina & Mauk 1999; Porrill & Dean 2007), and is consistent with experimental work suggesting that plasticity at the late-learning site for cerebellum-dependent learning may be heterosynaptic (Menzies *et al*. 2010; Pugh & Raman 2006). Second, we considered an internal feedback loop from the late-to the early-learning area of the circuit, in contrast to the feedforward architecture used in previous models. This internal feedback loop, which carries an efference copy of motor commands, has been suggested to be essential for producing the dynamics of individual eye movement responses (Lisberger & Sejnowski 1992; Lisberger 1994). We showed that this internal feedback loop also fundamentally changes the dynamics of memory consolidation, with stable consolidation requiring a circuit mechanism that drives plasticity so as to reset the activity of the early-learning area post-training (Fig. 6). This post-training reset provides a possible explanation for the previous suggestion, based on *in vivo* recordings (Lisberger 1994; Miles & Lisberger 1981), that the weights of the parallel fiber input pathway to Purkinje cells may paradoxically undergo potentiation in cases where the standard hypothesis of error-driven cerebellar cortical plasticity predicts that this pathway undergoes depression. Our work suggests one possible resolution to this controversy: the initial synaptic changes are governed by depression (Inoshita & Hirano 2018; Jang *et al*. 2020), but during consolidation the post-training reset mechanism reverses the decrease in weight, driving the weight to a larger steady state value relative to pre-training (Fig. 6*E,J* ; see Payne *et al*. 2024 for further discussion as well as an alternative solution). Third, we propose two biologically plausible circuit mechanisms that can perform this reset, suggesting new functional roles for elements of the cerebellar microcircuit in stabilizing a memory. The first mechanism, plasticity of the inhibitory feedforward inputs to the early-learning area, could be implemented by a Hebbian rule at the molecular layer interneuron (MLI)-to-Purkinje cell synapses (Vogels *et al*. 2011). Although these synapses are known to be plastic (Hirano 2018; Mapelli *et al*. 2015), weight changes were believed to be governed by climbing fiber-driven complex spikes in the postsynaptic Purkinje cell. Our model predicts that weight changes may instead be driven by the correlation of presynaptic MLI activity with Purkinje cell simple spikes. The second mechanism, inhibition of instructive signals by the late-learning area, could be implemented by an inhibitory pathway from the vestibular and cerebellar nuclei targeted by Purkinje cells to the inferior olive (Najac & Raman 2015; Uusisaari & Knöpfel 2011; see Herzfeld *et al*. 2020 and Kenyon *et al*. 1998 for related ideas), and might explain nonvisual climbing fiber responses that have been observed in the oculomotor cerebellum (Fanning *et al*. 2021; Winkelman & Frens 2006).

Learning and memory systems enable an organism to transform transient experiences into persistent effects. In working memory, neural integrator circuits accumulate and store transient input signals as persistent spiking activity. Here, we extend the concept of neural integration to the context of long-term memory by framing systems consolidation as a temporal integration process. Thus, this work suggests a unifying conceptual framework for describing computations underlying short- and long-term memory function.

## Acknowledgements

We thank Alireza Alemi, Hannah Payne, Friedemann Zenke, Daniel Fisher, Emre Aksay, Brian Wiltgen, Michale Fee, Ilana Witten, David Tank and Jay McClelland for helpful insight and comments. This work was supported by a Stanford Interdisciplinary Graduate Fellowship and a Stanford Center for Mind, Brain, Computation and Technology Traineeship to BJB, NIH grant R01 DC004154 to JLR, a esearch to Prevent Blindness grant to UC Davis Ophthalmology and NIH grant R01 EY021581 to MSG, and Simons Collaboration on the Global Brain grant 543031 and NIH grants R01 EY031972 and R01 NS072406 to JLR and MSG. Portions of the paper were developed from the thesis of BJB (Bhasin 2021). We assessed the gender and racial representation of authors we cite following the approach outlined in Dworkin *et al*. (2020), using open source code to evaluate the first names of the first and last authors (Zhou *et al*. 2022). Excluding self-citations to the first and last authors of our current paper, our references in the main text contain: 7.62% woman (first author)/woman (last author), 8.57% man/woman, 14.29% woman/man, and 69.52% man/man; and 12.99% author of color /author of color, 15.97% white author/author of color, 17.79% author of color/white author, and 53.26% white author/white author.

## Materials and methods

Code for simulations, analyses and producing plots for figures can be found at https://github.com/goldman-lab/consolidation-integration.

### Feedforward sensorimotor circuit model

Here we describe the feedforward model of Figures 2–4. The circuit model has parallel pathways that transform a graded, time-varying input into a response. In particular, we modeled the transformation of a graded, time-varying sensory (vestibular) input into an eye movement response and its modification by cerebellum-dependent learning. We modeled each node in the circuit with a single variable representing the average firing rate of a population of cells: mossy fibers, MF(*t*); parallel fibers (granule cell axons), PF(*t*); Purkinje cells, PC(*t*); medial vestibular nucleus neurons, MVN(*t*); and climbing fibers, CF(*t*) (Fig. 2*A*). Specific values of model parameters used in simulations were taken from the literature, where available, and are summarized in Table S1. For analysis, unless otherwise stated, all parameters are assumed to be nonnegative.

The firing rate of mossy fibers was modeled as a combination of a spontaneous baseline and a time-varying component encoding a sensory (vestibular) input driven by head motion 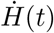,

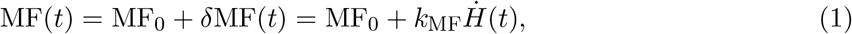

where we considered the encoding of the sensory input to be linear with sensitivity to head velocity *k*_MF_ (Lasker *et al*. 2008). Rather than explicitly model the mossy fiber to granule cell transformation, we considered the firing rate of parallel fibers to similarly depend on vestibular input linearly,

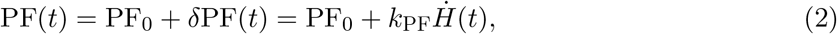

where *k*_PF_ was the firing rate sensitivity to head velocity.

Purkinje cells, in the early-learning area, were modeled as linearly combining direct excitatory input from parallel fibers and indirect inhibitory input via a parallel fiber-molecular layer interneuron pathway (Yamazaki *et al*. 2015),

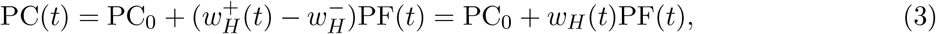

where 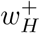 and 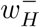 are the total excitatory and inhibitory weights onto the Purkinje cell, so the early-learning weight 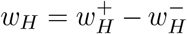 is the net weight of the parallel fiber pathway onto Purkinje cells. For simplicity, the inhibitory weight was not plastic in the model; the dynamics of learning are driven by the net weight. Equivalently, *w*_*H*_(*t*) can be interpreted simply as modeling the dynamics of the net (excitatory minus inhibitory) weight onto Purkinje cells. The effect of climbing fibers, which fire at very low rates (∼ 1 sp/s), on Purkinje cell output was not modeled here.

The medial vestibular nucleus, the late-learning area, was modeled as responding linearly to its excitatory mossy fiber input and inhibitory Purkinje cell input (Bagnall *et al*. 2008; Beraneck & Cullen 2007),

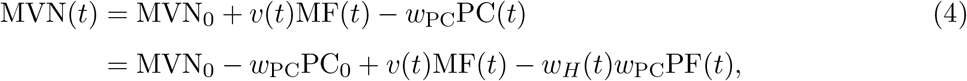

where *v* is the late-learning weight of the vestibular mossy fiber input to the medial vestibular nucleus, *w*_PC_ is the weight of the Purkinje cell input to the MVN, and MVN_0_ is an offset chosen so that the spontaneous MVN firing rate matched experimental values (57 sp/s) (Beraneck & Cullen 2007). The initial value of *v* before training was chosen so that the gain in the dark was 0.4 (Kimpo *et al*. 2014; Boyden & Raymond 2003).

Eye movement output, *Ė*, was modeled as being proportional to the deviation of medial vestibular nucleus neuron firing from its baseline (Yamazaki *et al*. 2015; Clopath *et al*. 2014),

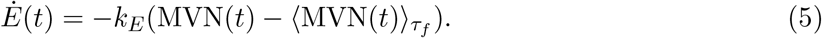

The baseline, 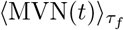, was calculated as an exponential (low-pass filtered) average over the recent past of MVN activity. For any time-varying quantity *x*(*t*), its exponential average ⟨*x*(*t*)⟩_*τ*_ over timescale *τ* is defined by

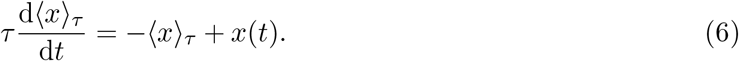

The timescale of the average *τ*_*f*_ = 0.017 h (1 min) is long relative to variations in the sensory input, but is much shorter than the timescales of plasticity (defined below), so that 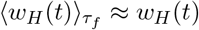 and 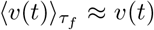 (Yamazaki *et al*. 2015). Throughout the paper, we assume that over the timescale *τ*_*f*_, the sensory input is zero on average,

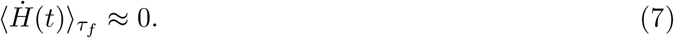

In the specific case of the bidirectional vestibular input to the oculomotor circuit, this corresponds to the time average of the rightward and leftward head movements being approximately equal.

The input-to-output gain of the circuit, *g*, is defined as the ratio of eye velocity output to head velocity input,

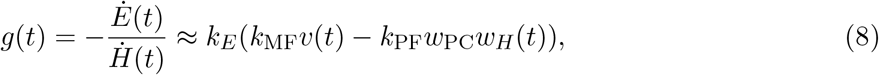

where the negative sign is present because eye movements are oppositely directed from head movements to keep the image of the world stable.

### Learning rules

Plasticity at the early-learning site was driven by feedback about behavioral errors, carried by the climbing fibers. Climbing fiber firing was represented by the sum of a spontaneous baseline rate and a time-varying component representing oculomotor errors,

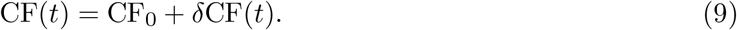

The time-varying component *δ*CF(*t*) encodes retinal slip, or a failure of eye movements to stabilize images on the retina, which occurs when the input-output gain of the circuit is incorrect. During training, the target output *Ė*^target^ of the circuit is a (negatively) scaled version of the vestibular input 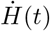 with gain *g*^target^,

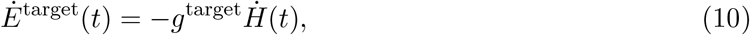

so, using Eq. (8), the retinal slip error is

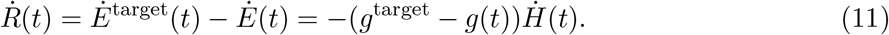

We model the time-varying component of the climbing fiber firing as

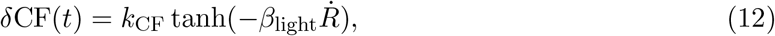

which is a saturating function of retinal slip. We chose the magnitude of the saturation to be *k*_CF_ = CF_0_ = 1 Hz, so that when retinal slip errors are large, climbing fiber firing saturates at a maximum of 2 sp/s and a minimum of 0 sp/s (Guo *et al*. 2014; Winkelman & Frens 2006). When retinal slip errors are relatively small (relative to 1*/β*_light_), *δ*CF(*t*) is approximately linearly related to negative retinal slip. Thus, if the eye movements are too small so that the gain of the response needs to be increased, the covariance between the climbing fiber and parallel fiber firing rates is positive.

Experimentally, coactivation of parallel fibers and climbing fibers causes associative long-term depression at the early-learning site, whereas activation of parallel fibers alone causes long-term potentiation (Raymond & Lisberger 1998; Suvrathan *et al*. 2016; Lev-Ram *et al*. 2002). Therefore, plasticity at the early-learning site was modeled as

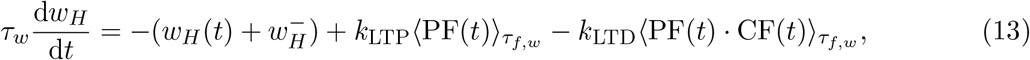

where the first term on the right is a passive decay, or forgetting, in the value of 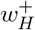 (Yamazaki *et al*. 2015; Clopath *et al*. 2014). We chose *k*_LTP_ and *k*_LTD_ so that the steady state value of 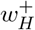 when not training was equal to 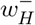, as required for stable consolidation (see SI Text, S1.1, for details). The time constant of plasticity, *τ*_*w*_, was chosen to be relatively fast (0.15 h = 9 min) during training (Boyden & Raymond 2003) and slower (5 h) during the post-training period (Jang *et al*. 2020; Cooke *et al*. 2004). The timescale of the averages over inputs, *τ*_*f,w*_, was chosen to be the same as *τ*_*f*_.

We considered two candidate learning rules for the late-learning site: a heterosynaptic and a Hebbian covariance-like rule. Heterosynaptic plasticity was modeled as

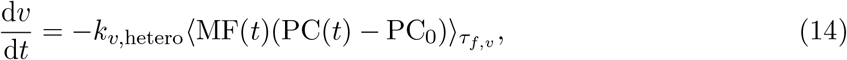

where an increase in *v* is driven by the anticorrelation between mossy fiber firing and fluctuations of Purkinje cell activity around baseline (Herzfeld *et al*. 2020; Clopath *et al*. 2014; Porrill & Dean 2007; Medina & Mauk 1999; Menzies *et al*. 2010; Pugh & Raman 2006). Hebbian plasticity was modeled as

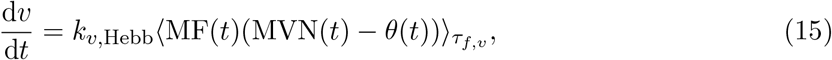

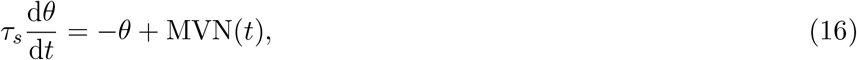

where the synaptic dynamics consist of the “readout” weight *v* and an internal sliding threshold variable *θ* that calculates a leaky average of postsynaptic activity (Yamazaki *et al*. 2015). The threshold is similar to those used in previous studies to counteract the runaway plasticity that would otherwise result from the inherent positive feedback driven by this kind of correlational rule (see SI Text, S4; Zenke *et al*. 2017; Miller & MacKay 1994; Bienenstock *et al*. 1982). For stability, the timescale of the sliding threshold was chosen to be fast relative to the rate of plasticity (see SI Text, S3, for details; Zenke *et al*. 2013).

### Simulation of oculomotor learning

We simulated oculomotor experiments with 0.5 h of training to increase the gain of the eye movement response, followed by a 23.5 h post-training period. During training, vestibular input was sinusoidal,

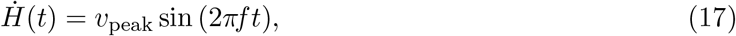

with peak velocity *v*_peak_ = 15 deg/s and rotational frequency *f* = 1 Hz, which is representative of experimental protocols (Jang *et al*. 2020; Kimpo *et al*. 2014; Nguyen-Vu *et al*. 2013; Boyden & Raymond 2003). In all simulations, we chose parameters such that the initial gain of the response was 0.4. In the simulations shown in Figure 2, we used a target gain value of *g*^target^ = 2, and chose 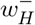, *k*_LTP_ and *k*_LTD_ so that during training, 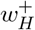 decreased by 51% (Jang *et al*. 2020) and the input-to-output gain increased by 30% from baseline (Kimpo *et al*. 2014; Boyden & Raymond 2003). To understand the stability of the dynamics during training, we also simulated a single 48 h training period to increase the gain to *g*^target^ = 1 (Fig. S1; see SI Text, S1.3). All other model parameters are shown in Table S1.

During the post-training period, we set 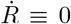. This models the animal being placed in the dark after training to eliminate feedback about oculomotor errors (Anzai *et al*. 2010; Shutoh *et al*. 2006; Kassardjian *et al*. 2005; Nagao & Kitazawa 2003; Boyden & Raymond 2003). We considered both the case in which vestibular input was not present post-training (as in Yamazaki *et al*. 2015), as well as the case in which it was present (as in Clopath *et al*. 2014). In the latter case, we used the same vestibular input as during training. For the heterosynaptic rule, the presence of post-training input had little effect on the change in *v*, unlike for the Hebbian rule (Fig. 5). The learning rates *k*_*v*,hetero_ and *k*_*v*,Hebb_ were picked so that approximately 75% of the increase in gain during the training period was consolidated (Boyden & Raymond 2003).

For the heterosynaptic rule, we studied the robustness of consolidated memories to perturbations in the early-learning site by adding random values to the early-learning weight *w*_*H*_ at regular intervals (Fig. 3). We initialized the weights to the same baseline as the training simulations and, every 10 minutes of simulation time, added a value chosen uniformly at random between −0.1 and 0.1 to *w*_*H*_. We simulated the effect of perturbations either in the absence of (Fig. 3*A–D*) or in the presence of (Fig. 3*E–H*) information about errors. For the simulations shown in Fig. 3*A* and *E*, we used the same parameters as for the simulation in Fig. 2 (Table S1), but with *τ*_*f,v*_ = *τ*_*f*_. For the simulations shown in Fig. 3*B* and *F*, we used a smaller learning rate, *k*_*v*,hetero_ = 6.95 × 10^*−*6^ (s/sp)^2^*/*h. For Fig. 3*F*, we simulated four consecutive days of training. We performed 250 runs of the perturbation simulations with the normal and smaller values of *k*_*v*,hetero_, with each simulation run for 24 hours of simulation time. The histograms in Fig. 3*C* were constructed from the endpoints of the 250 simulations in *A* and *B*, and we plotted the time course of the variances of the corresponding distributions (at the time point immediately before a perturbation) in Fig. 3*D*. Similarly to Fig. 3*C*, the histograms in *G* were generated from the final value of *v* across the 250 simulations in *E* and *F*, and *H* shows the time courses of the variance of *v* calculated in the same manner as for panel *D*.

All simulations were performed in Python by integrating the differential equations for the learning rules with the Radau solver (implemented in `solve_ivp` in the `scipy.integrate` package). Time steps in the simulation had a base unit of hours.

### Mitigation of stability-plasticity dilemma with two sites of learning

Here we outline some simple analyses we performed to understand how the stability-plasticity dilemma is addressed in the presence of noisy instructive signals driving learning in the circuit. We modeled learning across multiple training sessions where, during each session *k*, the target gain *g*^target^ = *ĝ*^(*k*)^ used to calculate retinal slip was randomly drawn from a normal distribution. Target gain values were drawn independently across training sessions. We built simplified versions of the model with one and two sites of plasticity, in which we ignore the exact time course of the weight dynamics and instead describe how the weights change discretely across training sessions (SI Text, S2; Fig. 4*B,F*). Using these simplified models, we calculated the mean squared error expected at the start and end of a typical training session as functions of the learning rates in the circuit (SI Text, S2.2; Fig. 4*C,E*). To build intuition for the difference between the one-site and two-site models more concretely, in Fig. 4*A* and *D* we also show the detailed time course of the gain and the corresponding squared error (retinal slip) of the full, unsimplified one- and two-site models for two subsequent example training sessions (see SI Text, S2.2, for details).

### Circuit model with internal feedback loop

Here we describe the expanded version of the circuit model that includes both a feedforward pathway carrying the sensory (vestibular) input to the early-learning area (Purkinje cells), as well as an internal feedback loop carrying the output from the late-learning area (MVN) back to the early-learning area (Fig. 6*A,F*). In the context of oculomotor learning, this internal feedback loop represents an efference copy of the output motor command. This expanded model has the same architecture as the Lisberger-Sejnowski model of oculomotor learning (Lisberger & Sejnowski 1992), but here we model the dynamics of the granule cell pathways as instantaneous. We show two candidate circuit mechanisms by which stable consolidation can be achieved when internal feedback is present.

The vestibular stimulus is carried by mossy fibers, described as before by Eq. (1), and by vestibular parallel fibers with firing rate

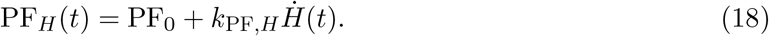

The efference copy i s carried through a different set of parallel fib ers with firing rate

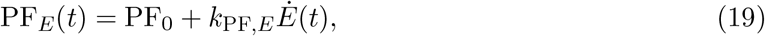

where the eye velocity output, Ė, is defined as before in Eq. (5). The firing rate of Purkinje cells is then a linear combination of vestibular and efference copy input, with a spontaneous baseline rate, i.e.,

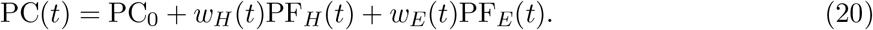

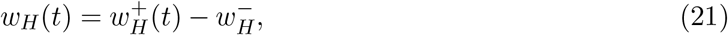

We assume that both parallel fiber pathways result in monosynaptic excitation and disynaptic inhibition onto the Purkinje cell, so that

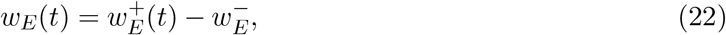

where 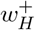 and 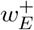 are the excitatory weights of the feedforward and feedback parallel fibers and 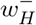 and 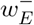 are the inhibitory weights of the interneurons in each pathway. Thus, as before, we can interpret *w*_*H*_ = 0 and *w*_*E*_ = 0 as balanced excitatory and inhibitory weights for each pathway.

The firing rate of MVN neurons is defined as before in the first line of Eq. (4), except we substitute the new definition of Purkinje cell activity from Eq. (20). The late-learning weight of the direct pathway is still called *v*. Then, following the same reasoning as the derivation leading to Eq. (8), we can calculate the eye velocity output as

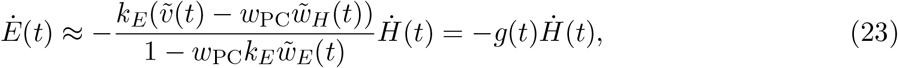

defining the prefactor in the second equality as the gain *g*(*t*), and where for simplicity we let

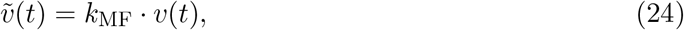

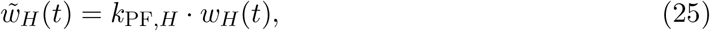

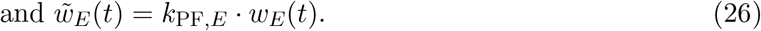

We use modified learning rules for the excitatory weights in the parallel fiber pathways, in which the weight values do not passively decay,

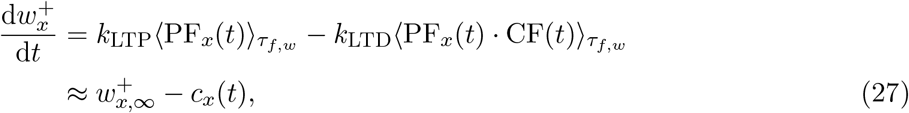

where *x* = *H* or *x* = *E* for the head pathway or efference copy pathway respectively, and 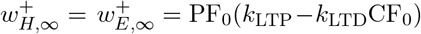 and 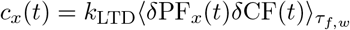. Here we assumed that *w*_*E*_ has a fixed nonnegative value for the purpose of visualizing the dynamics, but stable consolidation was also possible when *w*_*E*_ was plastic (Fig. S2; SI Text, S6). So that plasticity is completely controlled by the correlation between parallel fibers and climbing fibers, we assumed values of *k*_LTD_ and *k*_LTP_ such that 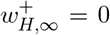. This condition can be interpreted as the plasticity due to spontaneous PF activity being offset by spontaneous CF activity (Kenyon *et al*. 1998).

For the direct pathway weight *v*, we used a modified version of the heterosynaptic rule in Eq. (14),

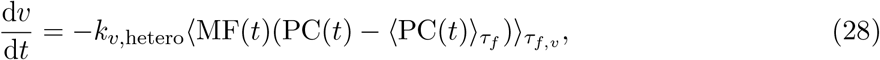

where the subtraction of the fixed Purkinje cell baseline has been replaced with a sliding threshold equal to the average PC activity over the recent past, similar to that used in the covariance-like Hebb rule, Eq. (15). The time constant of the sliding threshold was *τ*_*f*_, which was slow relative to variations in the sensory input but fast compared to the rates of plasticity.

As discussed in the main text, stable consolidation in the presence of a feedback loop requires that the circuit implement an active mechanism to reset early-learning activity post-training. We considered two such mechanisms. First, stable consolidation can be achieved if the feedforward inhibitory synapses onto the early-learning area 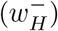 are plastic (Fig. 6*A*), governed by a Hebbian covariance-like rule,

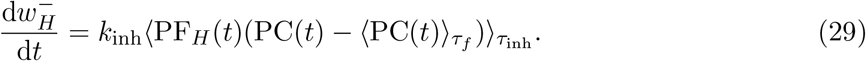

Second, stable consolidation can be achieved if, in addition to providing information about behavioral errors, the climbing fiber input at the early-learning site also serves to reset early-learning activity (Fig. 6*F*). We modeled climbing fiber activity as being driven by the instructive signal and inhibited by medial vestibular nucleus cells that receive only Purkinje cell inhibition, which corresponds to net excitation by PC activity,

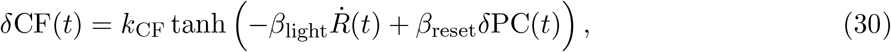

where 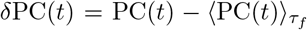. For both mechanisms, stable consolidation requires that the effective rate of plasticity of *v* be slower than that of *w*_*H*_ (SI Text, S5).

### Simulation of oculomotor learning in the model with feedback

We simulated the same oculomotor experiment as for the model without feedback (see *Simulation of oculomotor learning*), with differences in parameters specified in Tables S2 and S3. We used the same sensory input as for the feedforward model during both the training and post-training periods. We simulated the model with both the inhibitory plasticity (Fig. 6*A–E*, S3*A*) and inhibition of instructive signals (Fig. 6*F–J*, S3*B*) reset mechanism, assuming that *w*_*E*_ is not plastic. We used a target gain value of *g*^target^ = 2, and initial values 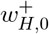 and *v*_0_ for 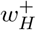 and *v*, respectively.

## SI Text

Below, we first provide mathematical analysis of the synaptic learning dynamics of the feed-forward circuit models with heterosynaptic (§S1, §S2) or Hebbian (§S3, §S4) plasticity rules at the late-learning site. We then analyze the learning dynamics in recurrent circuit models with either a fixed-strength (§S5) or plastic (§S6) internal feedback pathway.

## S1 Analysis of synaptic learning dynamics in the feedforward model

To understand the conditions for successful consolidation in the model, we examined the dynamics of learning analytically. To simplify our analysis, we consider timescales that are relatively long compared to variation in the sensory input, so that the input is zero on average, Eq. (7), and the variability of the sensory input, as measured by the time average of the squared input, is constant,

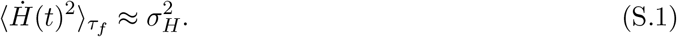

### S1.1 Post-training dynamics and conditions for stable consolidation

Using these assumptions, the dynamics of the early-learning site can be described by simplifying Eq. (13) to

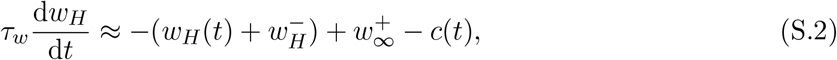

where 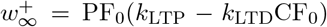 depends only on the baseline firing rates of inputs to the Purkinje cell, and 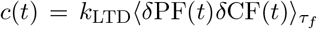 is proportional to the covariance of the parallel fiber-mediated sensory input and climbing fiber feedback about errors. When the circuit is producing behavioral output without error (as we assume is the case before training), or during the post-training period, when there is no feedback about errors, *c*(*t*) ≡ 0. As a result, *w*_*H*_ tends toward a baseline value of 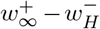. If *c*(*t*) *>* 0 during training, as was the case in our simulations, *w*_*H*_ decreases from this baseline, increasing the input-output gain of the circuit at the end of the training period. On the other hand, eye movements that are too large would result in *c*(*t*) *<* 0, causing an increase in *w*_*H*_ and a decrease in the gain.

At the late-learning site, the heterosynaptic rule, Eq. (14), can be simplified to

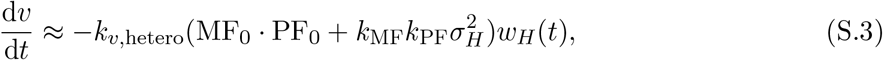

where we assume that the timescale of plasticity of *w*_*H*_ is significantly longer than *τ*_*f*_ (the timescale of correlations in neural activity to which the learning rule is sensitive) and therefore the value of *w*_*H*_ is effectively uncorrelated with the sensory input. From the form of Eq. (S.3), the late-learning weight *v* is proportional to the temporal integral of the early-learning weight *w*_*H*_.

We can then understand the dynamics of learning as follows. Initially, when errors are large and correlated with the sensory input, because of the saturation in the CF response, we have

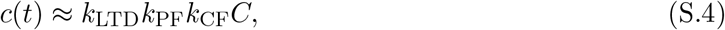

where

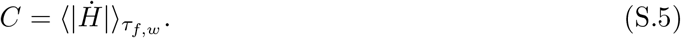

For sinusoidal sensory input with period of oscillation much shorter than *τ*_*f,w*_, *C* ≈ 2*/π* ·*v*_peak_, which is constant for a fixed choice o f p eak i nput a mplitude. D uring t raining, w hen *c* (*t*) ≠ 0, *w* _*H*_ tends toward a value that is smaller than baseline by an amount proportional to *C* (Fig. S1*A,B*). This saturation can be understood as long-term depression driven by error being balanced by the decay in *w*_*H*_ to baseline. As *v* starts to integrate, this further decreases the magnitude of error. When errors are decreased enough such that *c*(*t*) is approximately linear in terms of error magnitude,

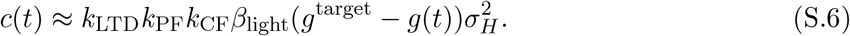

In this regime, there is a stable fixed point of the weights that corresponds to the target input-to-output gain (Fig. S1*B,C* ; see below and §S1.3).

After training ends, *w*_*H*_ returns to baseline. *v* must also go to a steady state in order for the system to be stable. From Eq. (S.3), this occurs as long as *w*_*H*_ has a baseline value of 0, which from Eq. (S.2) requires that

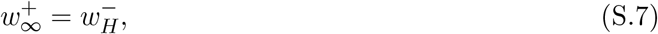

that is, when at steady state the excitatory and inhibitory weights of the sensory input to the early-learning site are balanced. As a result, there is a continuum of steady states of the system in the post-training period, corresponding to a continuum of input-to-output gains of the circuit, since any weight configuration where *w*_*H*_ = 0 is a fixed point of the dynamics, which can be represented as a line in *w*_*H*_-*v* space (Fig. 2*E*). We note that the stable fixed points of the system during training, as determined by the target gain *g*^target^, also lie along the same line (Fig. S1*C*). When the steady-state excitation-inhibition balance condition, Eq. (S.7), is met, any initial set of weights will evolve toward some fixed point on the line *w*_*H*_ = 0. Therefore, *w*_*H*_ = 0 is furthermore a line attractor. This can be visualized by plotting a flow field in *w*_*H*_-*v* space (Fig. 2*D,E*), where a vector describing the direction of a trajectory at any point is given by **x**(*w*_*H*_, *v*) = [d*w*_*H*_*/*d*t*, d*v/*d*t*]. The overall dynamics of learning and consolidation can thus be put succinctly: oculomotor errors drive a change Δ*w*_*H*_ in the early-learning weight that is temporally integrated into a persistent change Δ*v* in the late-learning weight, as the change in *w*_*H*_ decays away. The need to transform a negative Δ*w*_*H*_ into a positive Δ*v* explains why plasticity at *v* is driven by the *anti*correlation between mossy fiber and Purkinje cell fluctuations in Eq. (14).

A similar analysis for the Hebbian covariance-like rule shows that a line attractor may exist when there is no variability in the sensory input, 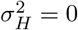 (Fig. 5*A,B*, light lines), but that the introduction of variability collapses the line attractor into a single unstable fixed point, which would not allow the circuit to learn an analog input-to-output gain (Fig. 5*A,B*, dark lines; §S3). As discussed in the main text (*Implications for plasticity rules*), simple implementations of classic homeostatic Hebbian learning rules that fix the norm of the weight(s) onto the neuron (Miller & MacKay 1994; Oja 1982) or attempt to maintain postsynaptic firing rate at a target value (Yger & Gilson 2015; Bienenstock *et al*. 1982) also failed to support a continuum of stable weight configurations in the model (see §S4). Our analysis found that a modified Hebbian covariance rule containing a decay could stably support a continuum of steady states even for 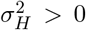, but this required meeting a fine-tuning condition that depends on the value of 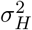 (see §S3).

### S1.2 Flow field analysis and diffusive drift

Using the flow field, we can understand how the learning rates at the early- and late-learning sites control the amount of consolidation after a period of training. We assume that the circuit starts in a consolidated state before training, so that during the training period *w*_*H*_ changes from its baseline value of 0 by an amount Δ*w*_*H*_. We also assume that *v* does not change much during the training period. From Eq. (S.2) and Eq. (S.3), the slope of the vectors **x**(*w*_*H*_, *v*) describing the flow during the post-training period is

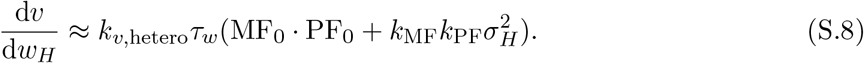

In other words, geometrically, the weight configuration moves approximately along a line with slope given by Eq. (S.8), so that as *w*_*H*_ returns to 0, the change in *v* during the post-training period is given by Δ*v* ≈ −d*v/*d*w*_*H*_ · Δ*w*_*H*_. Since the slope is proportional to *k*_*v*,hetero_*τ*_*w*_, a learning rate *k*_*v*,hetero_ at *v* that is much smaller than the learning rate 1*/τ*_*w*_ at *w*_*H*_ would correspond to slow consolidation, but also a corresponding insensitivity of *v* to noisy fluctuations in *w*_*H*_ during the post-training period (Fig. 3).

For the circuit to completely consolidate the changes made at the early-learning site during the training period—i.e., preserve the value of the gain *g*_train_ achieved during training—trajectories need to follow the line in synaptic weight space corresponding to a constant gain (Eq. (8), with *g*(*t*) ≡ *g*_train_). This is achieved if the slope of the flow field is close to that of the constant gain line,

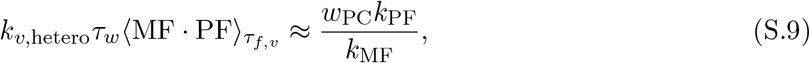

(see Fig. 2*D*, dashed line). In general, we can write the fraction *p* of the gain change induced during training that is consolidated post-training as

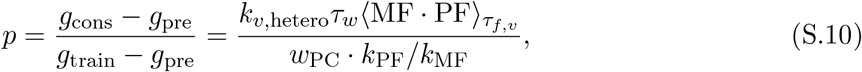

where *g*_pre_ is the gain before training, *g*_train_ is the gain immediately after training, and *g*_cons_ is the gain after consolidation. Complete consolidation (i.e., Eq. (S.9)) corresponds to *p* = 1.

As noted in the main text, because *v* is integrating, noise is also accumulated. We examined this by perturbing *w*_*H*_ in the absence of training signals from feedback about errors (see Materials and Methods, *Simulation of oculomotor learning*). Extending the current analysis to the case of random perturbations, we can show that, for a sequence of perturbations in *w*_*H*_ that are uniformly distributed over [−*α, α*], the variance across simulations of the value of *v* after the *k*th perturbation will be

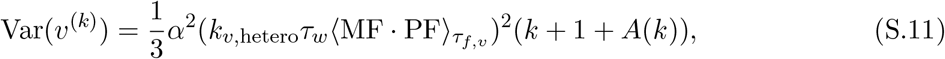

where *A*(*k*) is an exponentially decaying function of the number of perturbations and the time between perturbations (see §S1.4 for definition and derivation). Thus, the variance is asymptotically linear in *k* and the behavior of *v* is diffusive on long timescales. This is the key mathematical observation underlying the “speed-accuracy” tradeoff shown in Fig. 3.

The slope of the flow field is also related to the variability of sensory input 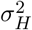. In this way, an input with larger variability would increase the rate of plasticity at *v*. However, this contribution to the rate of plasticity due to stimulus-driven variability is much smaller than that due to the high baseline firing rates of the mossy fibers MF_0_ and parallel fibers PF_0_ (not shown; see Eq. (S.8)) given the parameters we used, which were chosen to match experimental values.

### S1.3 Stability during training

During training, we assume that the climbing fiber modulation *δ*CF(*t*) is modeled by Eq. (12), i.e., it is a saturating function of the oculomotor (retinal slip) error 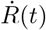 (Eq. (11)). Then, there is a fixed point of the weight dynamics corresponding to the point at which the measured gain of the system is *g* = *g*^target^,

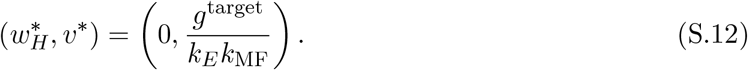

The dynamics of trajectories leading to a steady state can be separated into two regimes. Initially, the amplitude of oculomotor errors is large, so that *δ*CF saturates, and correspondingly so does the parallel fiber-climbing fiber covariance *c*(*t*). As a result, the weight *w*_*H*_ tends toward the value defined by Eq. (S.4). From the simplified form of the heterosynaptic plasticity rule for *v*, Eq. (S.3), this leads to initially linear growth in *v*. As *v* increases, eventually the amplitude of oculomotor errors will decrease, such that *δ*CF will be a linear function of 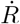, and from Eq. (S.6), *c*(*t*) will be a linear function of *w*_*H*_ and *v*, such that

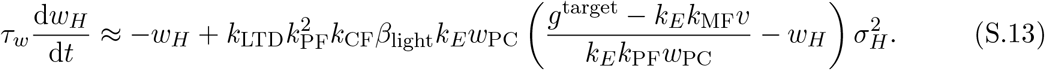

Thus, in the linear regime, the dynamics have eigenvalues

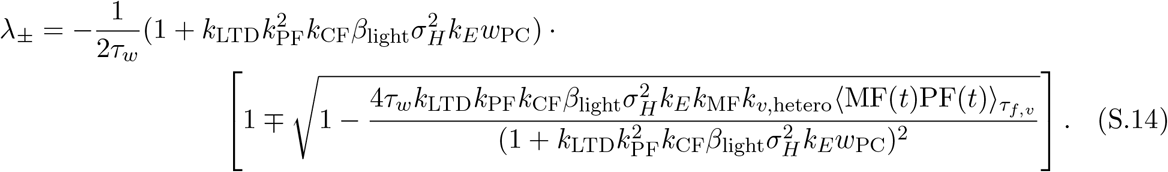

Since all of the parameters are strictly positive (during training,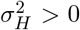), the steady state in Eq. (S.12) is a stable fixed point of the dynamics.

The approach to the fixed point could involve decaying oscillations in the values of *w*_*H*_ and *v* around the steady state. Oscillations do not occur if

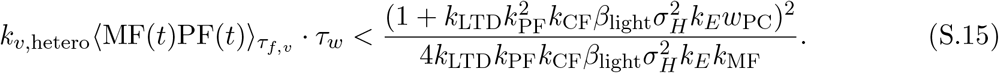

The above inequality places a constraint on the ratio of the rate of plasticity at *v* relative to the intrinsic rate of plasticity (1*/τ*_*w*_) at *w*_*H*_ to avoid oscillations.

The above analysis shows that, even when the size of climbing fiber responses decreases across learning as a result of decreased errors in the output, stable consolidation is still achieved. Learning at the early-learning site decreases in amplitude as consolidation occurs at the late-learning site until the system reaches desired performance, at which point the early-learning area is no longer required (Fig. S1).

### S1.4 Analysis of drift without error feedback

We noted above that consolidation could be understood as temporal integration at the late-learning site *v* of weight changes initially induced in the early-learning weight *w*_*H*_. Drawing on other work on integrators, we showed that periodically applied random perturbations of *w*_*H*_ led to the value of *v* random-walking over time (Fig. 3*A–D*). Here, we derive analytically a formula for the variance across sample paths of the late-learning weight *v* as a function of time and the learning rate at *v*. We start from the analysis in §S1.1 above. As in the simulation of Fig. 3*A–D* (see Materials and Methods, *Simulation of oculomotor learning*), we assumed that the value of *w*_*H*_ was perturbed at a regular interval *T*, with each perturbation *k* given by an i.i.d. random value 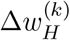 drawn uniformly over the interval [−*α, α*]. After the perturbation, the synaptic dynamics cause *w*_*H*_ to decay toward a baseline value of zero. To model the effect of each perturbation, we assume the value of *w*_*H*_ immediately following the *k*th perturbation is given by

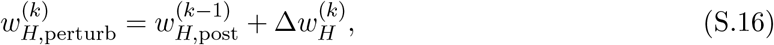

where the value of *w*_*H*_ just prior to the *k*th perturbation is given by

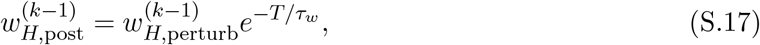

which models the decay in *w*_*H*_ between perturbations. We can simplify this recurrence relation as

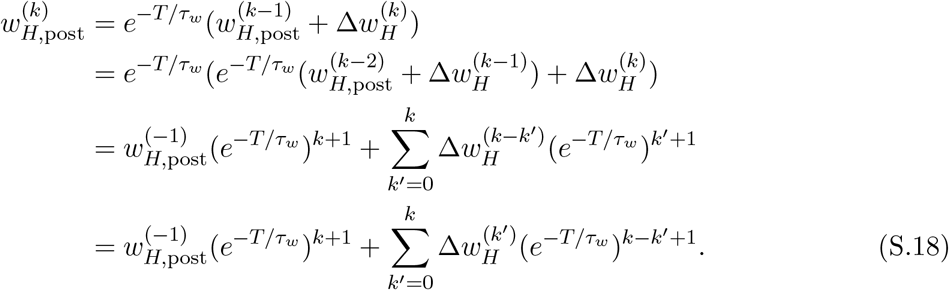

We assume that the system starts in a stable steady state, i.e.,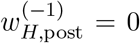. The change during each post-perturbation period *k* ≥ 0 is thus

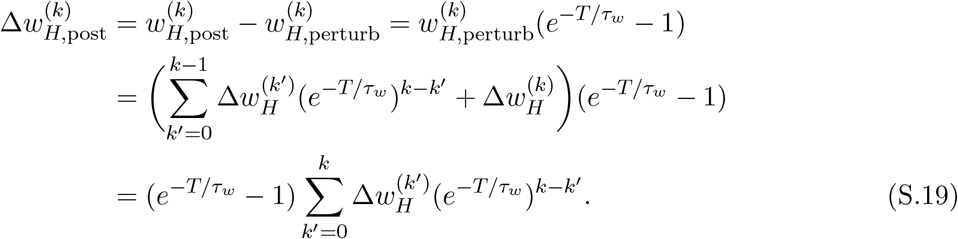

Then, from Eq. (S.8), the change in *v* during the *k*th perturbation period can be approximated as

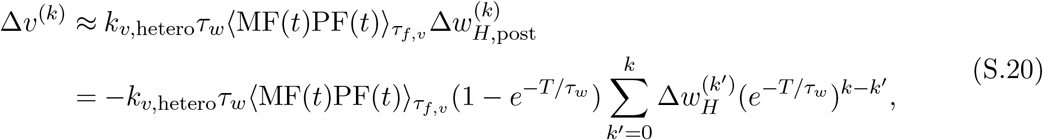

using the relationship in Eq. (S.19).

We can calculate the overall value of *v* after the *k*th perturbation as

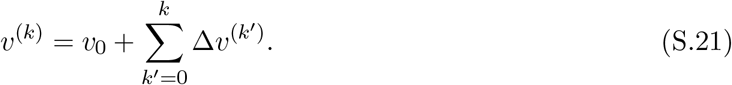

This has variance

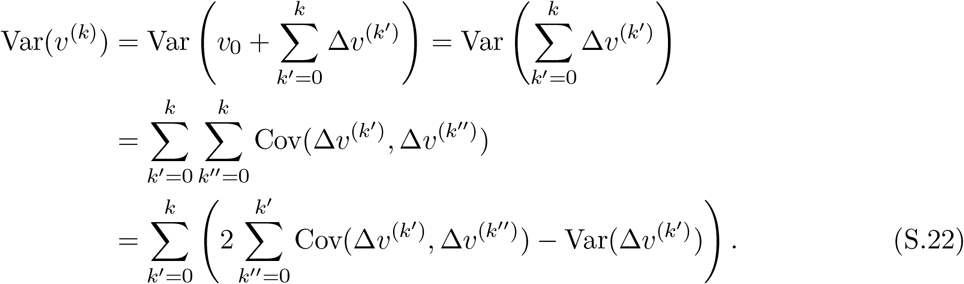

Thus, to calculate the variance in *v*^(*k*)^, we need to calculate the variance of Δ*v*^(*k*)^ and the covariance between Δ*v*^(*k*)^ and Δ*v*^(*j*)^. For convenience, define

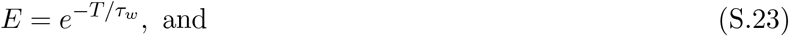

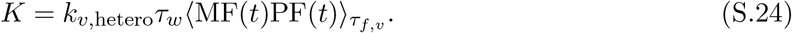

We can write the covariance between the change in *v* due to perturbations *j* and *k* as

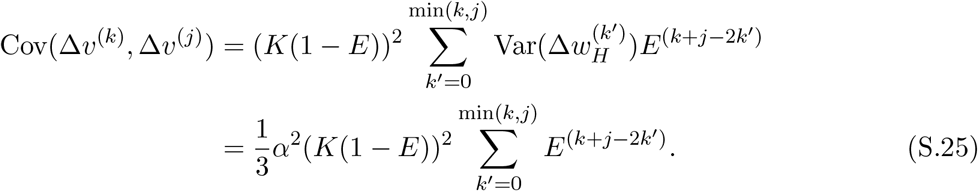

where, because 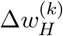 is uniformly distributed over [−*α, α*],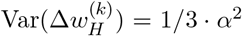. The variance of Δ*v*^(*k*)^ can be found by taking *j* = *k* in Eq. (S.25),

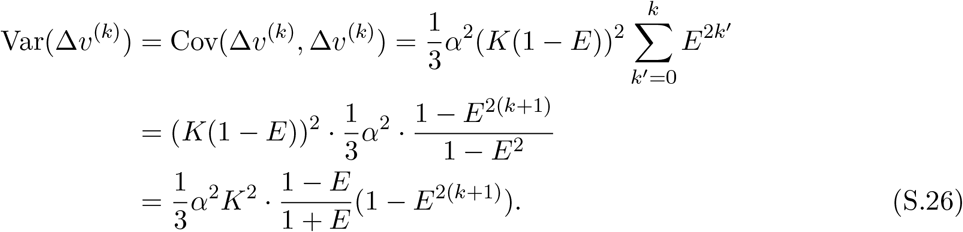

We can then calculate the sum

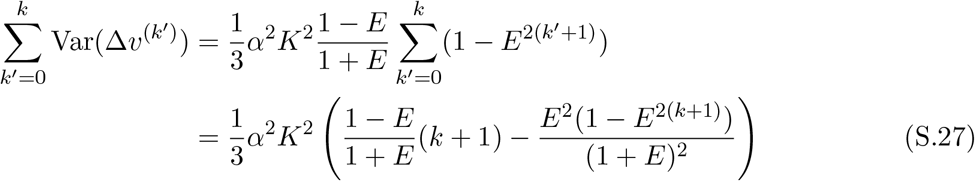

Finally, from Eq. (S.25), we can also calculate the sum of covariances

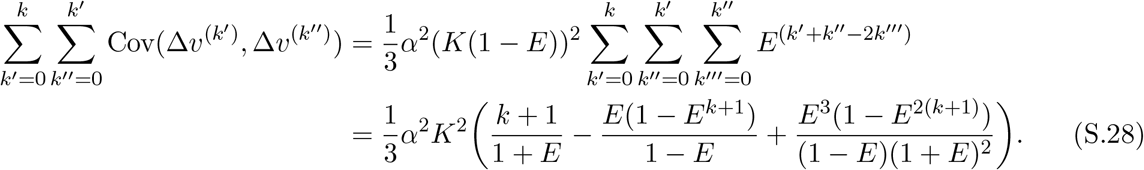

Putting these results together, from Eq. (S.22), the variance of *v*^(*k*)^ is

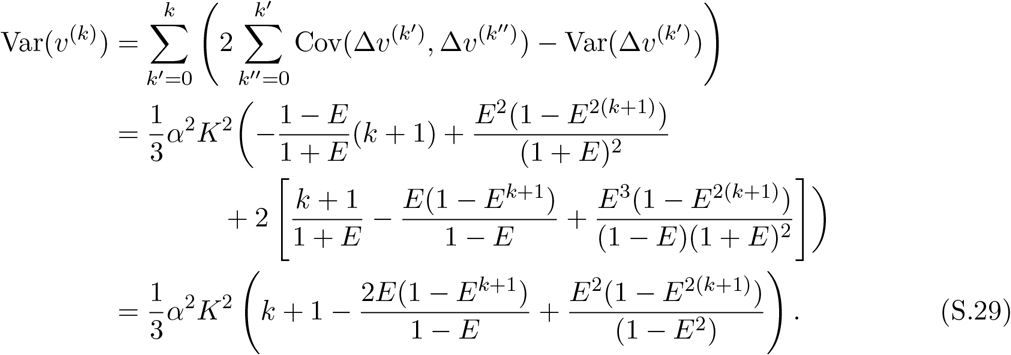

That is, the term *A*(*k*) in Eq. (S.11) is given by

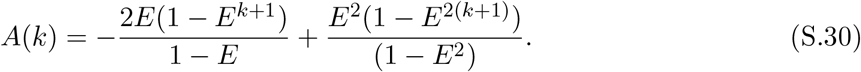

## S2 Mitigation of stability-plasticity dilemma with two sites of learning

To understand how consolidation helps to mitigate the stability-plasticity dilemma, here we build and analyze simplified discrete models of the weight dynamics. In the first section below, we show that consolidation, as an integration process, causes the circuit to effectively average over the noise in the instructive signal. In the following section, we show that averaging allows the circuit to mitigate a stability-plasticity tradeoff that would occur if the circuit only had one site of plasticity.

### S2.1 Consolidation averages over variability in instructive signals

We begin by considering a simplified version of the full feedforward model of Figs. 2 and 3. We assume that *w*_*H*_ returns to a baseline value of zero after each training period, and that *v* perfectly integrates. During a training session *k* ≥ 0, the model receives a retinal slip error signal corresponding to a target gain value *g*^target^ = *ĝ* ^(*k*)^ that is randomly drawn from a normal distribution, and learns a fraction *q* of the change in *w*_*H*_ that would fully minimize the error. From Eq. (8) and Eq. (11), this can be written as

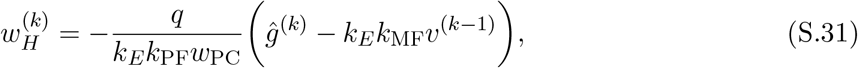

so that the output of the circuit after training has measured gain

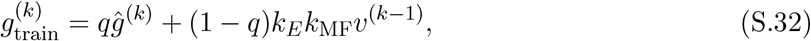

where we assume that the change in *v* during training is negligible, and that 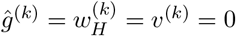 for all *k <* 0. During the subsequent post-training period, the model consolidates a fraction *p*_*w*_ of the learned weight change in *w*_*H*_ into *v*,

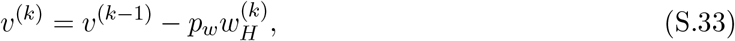

and *w*_*H*_ returns to a baseline value of zero, so the measured gain after consolidation is

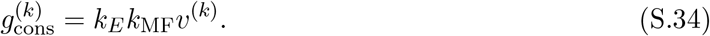

Then, the relative gain change consolidated is

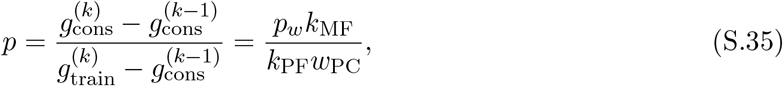

and we can rewrite the change in *v* after consolidation and the measured gain as

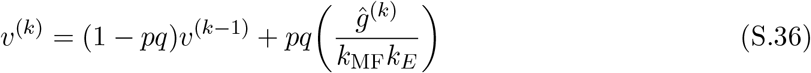

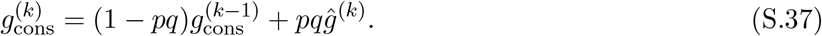

To solve the recurrence relation above, we take the unilateral *z*-transform of Eq. (S.37):

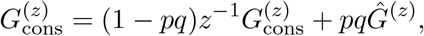

where the capital letters represent *z*-transformed functions. We can rearrange this as

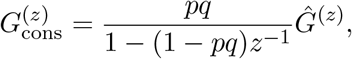

and take the inverse *z*-transform to find

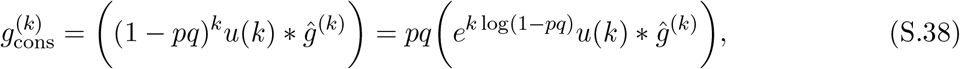

where the star operator represents discrete convolution and the discrete unit step function is defined to be *u*(*k*) = 1 for *k* ≥ 0 and 0 for *k <* 0. Intuitively, we can understand the measured gain after consolidation as taking a leaky average of the noisy target gain signal *ĝ*^(*k*)^ over the recent past with time constant (in terms of training sessions) approximately −1*/* log(1 − *pq*).

Given that *ĝ*^(*k*)^ is a random variable, we can also understand Eq. (S.37) as an autoregressive AR(1) process, whose stationary distribution has mean and distribution

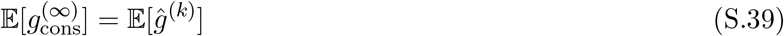

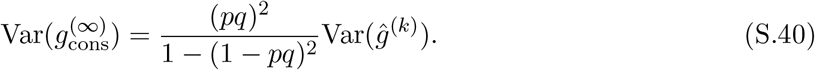

That is, the stationary distribution of the measured gain after consolidation has mean equal to the true mean of the noisy target gain distribution, and variance that is an increasing function of the consolidation fraction *p*.

From Eq. (S.38) we can see that a system that consolidates a large fraction of what is learned during the training period, *p* → 1, averages over a relatively small amount of the recent past—for fast early-learning, *q* = 1, the time constant approaches zero as *p* → 1. Furthermore, if the mean of the noisy target gain distribution changes, the approach of the consolidated gain to the new true mean will be fast. However, as a result, the variance of the measured gain across training sessions will have a relatively large variance, from Eq. (S.40). On the other hand, a system that consolidates slowly (*p* → 0) averages over a longer period, reaching the true mean slowly but with a lower stationary variance.

These features of the model are illustrated in Figure 4*F*. We plotted the value of the consolidated gain 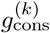 resulting from simulating the simplified model, Eq. (S.31) and Eq. (S.33), with *p* = 0.1 (“slow consolidation”, magenta) or *p* = 0.75 (“fast consolidation”, cyan), in response to the same set of target gain values. Initial values of *w*_*H*_ and *v* and all other parameters are the same as for the full model above (Table S1). For the first 50 training sessions, target gain values were drawn from a distribution with mean 0.4 and standard deviation 0.1, and for the subsequent 150 sessions, from a distribution with the same standard deviation but with mean 2.

### S2.2 Comparing a one-site and two-site model

To see the advantage of the averaging effect described above, we consider a one site version of the simplified model (effectively, the late-learning weight is constant), with weight *w*_*H*,1_ and no consolidation. From Eq. (8), the measured gain of this circuit after each training period *k* is then given by

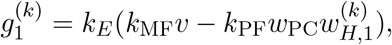

where *v* is fixed at the initial value *v*_0_. During each training period, the model again learns a fraction *q* of the total weight change in *w*_*H*,1_ required to minimize error,

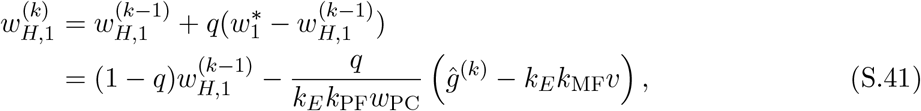

or equivalently in terms of the measured gain,

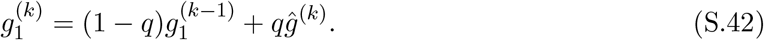

Note that the weight update, Eq. (S.41), is equivalent to the weight update for the two-site model, Eq. (S.31), if the weight were reset post-training, i.e., if 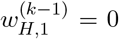 after consolidation. That is, just as for the two site model, the measured gain of the system is an AR(1) process representing a leaky average of the target gain signal with time constant −1*/* log(1 − *q*). For a sequence of target gain values drawn from a normal distribution, the stationary distribution of the measured gain of the circuit will have mean equal to the true mean and variance

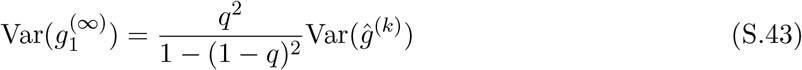

which is minimized as *q* becomes small (“*w*_*H*_ slow” in Fig. 4*A–C*). For such a small value of *q*, the measured gain will remain relatively stable across training sessions, but the circuit will be unable to quickly correct errors. That is, after each training session, the mean squared error of the one-site model in the stationary limit will be

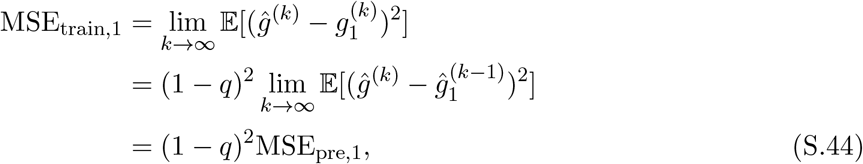

where MSE_pre,1_ is the mean squared error at the start of each training session,

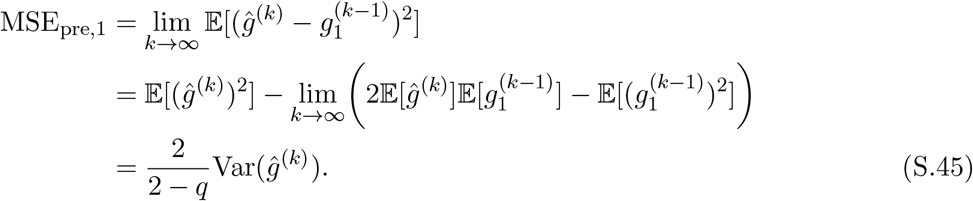

That is, after training to adapt to a target gain experienced during training session *k* − 1, MSE_pre,1_ represents the mean squared error expected at the start of training session *k*, when a new target gain value will be drawn. From Eq. (S.44) and Eq. (S.45), we see that the average error after training is a decreasing function of *q*, whereas the average error at the start of training is an increasing function of *q* (for *q <* 2). Hence, there is a tradeoff between the ability of the circuit to correct errors quickly during a training session and the size of the expected future error given a noisy target gain.

Repeating this calculation for the model with two sites of learning, we have

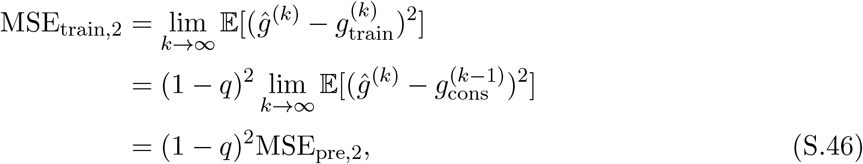

and

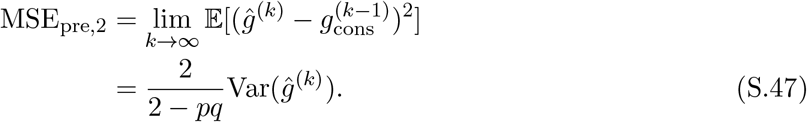

Thus, the average error after training can be reduced by fast learning at the early-learning site (*q* → 1), and the average error at the start of training can be simultaneously reduced by slow learning at the late-learning site (*p* → 0).

In Figure 4*C,E*, we plotted the one-site tradeoff curve as functions of the early site learning rate *q* (“*w*_*H*_ slow” and “*w*_*H*_ fast” arrows), with MSE_pre,1_ on the *x*-axis and MSE_train,1_ on the *y*-axis (black line in Fig. 4*C* and grey line in *E*), normalized by plotting in units of the variance of the target gain distribution. For the two-site model (Fig. 4*E*), we plotted the normalized MSE_pre,2_ on the *x*-axis and normalized MSE_train,2_ on the *y*-axis for *p* = 0.1 (“*v* slow”, magenta) and *p* = 0.75 (“*v* fast”, cyan), while again varying *q*. In Figure 4*B*, we plotted the post-training gain 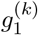 from a simulation of the simple one-site model with *q* = 0.1 (“slow learning”, black) or *q* = 0.75 (“fast learning”, grey) over 200 training sessions, where the target gain value for each session was independently drawn from a normal distribution with mean 0.4 and standard deviation 0.1. The values of MSE_pre,1_ and MSE_train,1_ corresponding to these values of *q*, calculated from Eq. (S.44) and Eq. (S.45), are plotted in Figure 4*C* with circles.

To build intuition for the difference between the one-site and two-site models more concretely, in Figure 4*A,D* we show the time course of the gain and the corresponding squared error (retinal slip) for two subsequent example training sessions. Figure 4*D* was generated by simulating the full two-site model with weight dynamics controlled by Eq. (S.2) and Eq. (14). We set the learning rate at *v* to *k*_*v*,hetero_ = 1.29 × 10^*−*6^(s/sp)^2^*/*h, which resulted in a fraction of learning consolidated *p* ≈ 0.13 for the *k*th and *p* ≈ 0.27 for the (*k* + 1)th sessions. All other parameters were the same as those in Table S1, and the initial value of *v* was set so that the system would have an initial gain of 0.4, as above. To similarly simulate the time course of learning in the one-site model (Fig. 4*A*), we modified the full two-site model by removing learning at *v*, and the learning at *w*_*H*_ was governed by

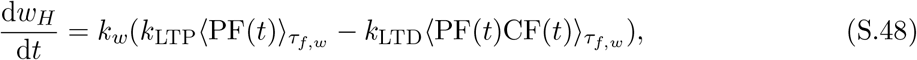

where we set the learning rate parameter *k*_*w*_ = 0.191 h^*−*1^ (light colors) or *k*_*w*_ = 1.97 h^*−*1^ (dark colors), so that for the two training sessions shown the fraction learned was *q* ≈ 0.10 for the *k*th session and *q* ≈ 0.11 for the (*k* + 1)th session (light colors), or *q* ≈ 0.75 for the *k*th and *q* ≈ 0.80 for the (*k* + 1)th sessions (dark colors). Unlike the two-site model, we set *k*_LTP_ = 0.648 s/sp, so that outside of the training period, d*w/*d*t* ≈ 0. The model used the same initial values, as well as all other parameters not specified here, as the two-site model for *w*_*H*_ and *v*. For both models, we simulated two periods of 0.5 h training followed by 11.5 h post-training without feedback about behavioral errors (post-training periods clipped for illustrative purposes).

## S3 Consolidation with the Hebbian rule is not robust to noisy input

The failure of the stabilized Hebbian rule, Eq. (15), to successfully consolidate memory from the early-to late-learning site in the presence of variability in the sensory input (Fig. 5) can be understood through an analysis similar to the one performed for the heterosynaptic rule (§S1). We again assume that the mean of the sensory input is approximately zero and that the variability 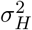 is constant over the timescales *τ*_*f,v*_ and *τ*_*s*_. We also assume, as before, that the timescale over which the weights change is much slower than *τ*_*f,v*_. Using that the threshold *θ*(*t*) is the exponential average of MVN(*t*), we have

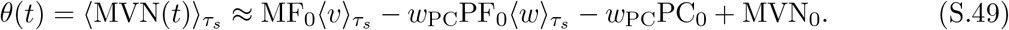

Then, substituting this and the definition of MVN(*t*), Eq. (4), into the learning rule, Eq. (15) becomes

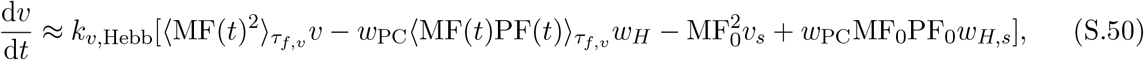

where 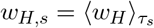 and 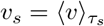.

In the post-training period, when *c* → 0, we have that 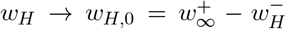 approaches this steady state, so will the exponential average, i.e., *w*_*H,s*_ → *w*_*H*,0_ (see Eq. (6)), so we can use this quasi-steady state assumption on *w*_*H*_ to simplify Eq. (S.50) further to

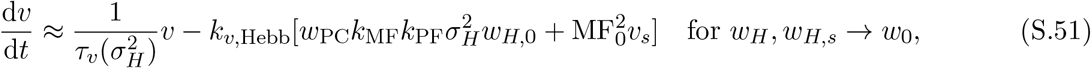

where the time constant *τ*_*v*_ is a function of the variability of the sensory input, 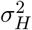,

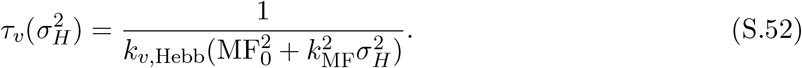

In the case that there is no variability in the sensory input, 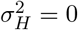, we have that the dynamics of learning are given by

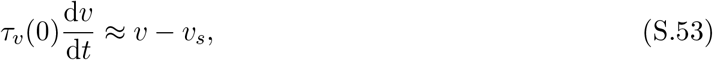

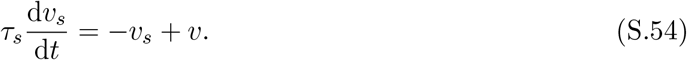

Here, the first eigenvalue of the system is 0, with corresponding eigenvector [1, 1] so that any state along the line *v* = *v*_*s*_ is a fixed point of the system. The second eigenvalue of the dynamics is

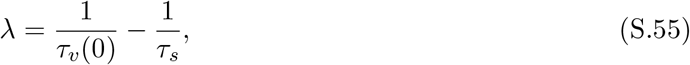

which is decaying if *τ*_*s*_ *< τ*_*v*_(0), i.e., the timescale of the sliding average is faster than the timescale of plasticity (cf. Zenke *et al*. 2013). Thus, the synaptic learning dynamics contain a line attractor in the space of *v* and the internal variable *v*_*s*_ that does not depend on *w*_0_. Geometrically, when 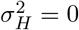, the nullclines for the dynamics of *v* and *v*_*s*_ overlap, creating the line attractor.

If there is variability in the sensory input, 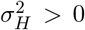, the nullclines no longer exactly overlap; there is now a single fixed point 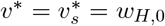. This fixed point is unstable (it is a saddle point of the dynamics), with eigenvalues

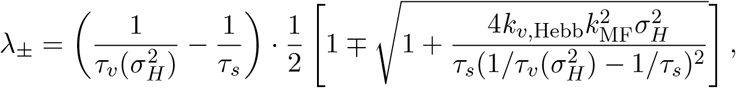

implying that one of the eigenvalues will always be positive.

In principle, the instability can be rectified by adding an additional decay term −*εv* to the learning rule, so that Eq. (15) is

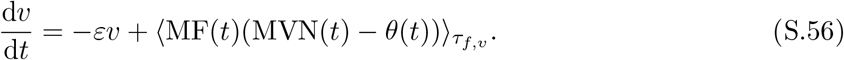

If we write the decay rate as 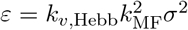 in terms of a new variable *σ*^2^, we can simplify this equation (as we did for Eq. (S.51)) to

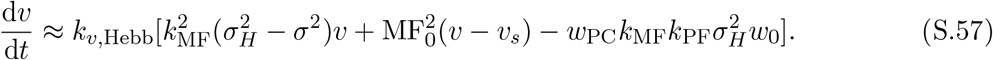

If *σ*^2^ is equal to 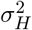, the variability of the vestibular input in some period, and if *w*_*H*,0_ = 0, then the system will again be described by the dynamics given by Eq. (S.53) and Eq. (S.54). That is, *v* = *v*_*s*_ will still be a line attractor. If *σ*^2^ is not exactly equal to 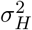, then the weight will either grow unstably or decay to zero, depending on whether 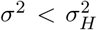 or 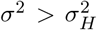, respectively. This can be seen by assuming the threshold *θ* moves instantaneously, so that *v*_*s*_ ≈ *v*, and noting that Eq. (S.57) becomes either exponential growth or decay, with time constant inversely proportional to the degree of mistuning. So, for successful systems consolidation in a circuit using this modified version of the Hebbian rule, the learning rule must “know” or otherwise measure the variability of the input 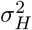. While a learning rule that accounts for the natural statistics of inputs presents an interesting possibility, we note that it may be challenging to implement biologically.

## S4 Traditional homeostatic Hebbian rules do not readily support consolidation of analog memories

It has long been recognized that the basic form of the Hebbian learning rule generally leads to unstable positive feedback (Miller & MacKay 1994; Zenke *et al*. 2017; Abbott & Nelson 2000), leading to the development of rules which contain homeostatic mechanisms that attempt to counter this instability. The most well-known classes of such homeostatic mechanisms include weight normalization, in which weights are adjusted so that the weight vector for a neuron maintains a given length (von der Malsburg 1973; Oja 1982; Miller & MacKay 1994), and firing rate-target homeostasis methods, in which weights are adjusted so that the mean postsynaptic firing rate is maintained close to a specified target. The latter class of mechanisms includes those commonly called synaptic scaling (Turrigiano 2008; Yger & Gilson 2015; Tetzlaff *et al*. 2011). In order to stably consolidate analog memories, the learning rule at the late-learning site must be able to support a continuum of stable weight configurations. For the circuit model we developed here, we show below that Oja’s rule and the BCM rule, prominent examples of weight normalization and firing rate-target homeostasis rules, respectively, cannot support such a stable weight continuum.

First, we examine Oja’s rule, a form of weight normalization (Oja 1982). We will consider the Hebbian piece in a “covariance” form,

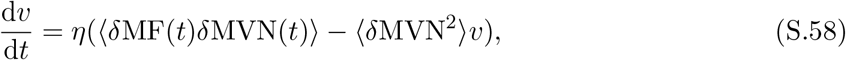

where for simplicity, *δ*MF(*t*) and *δ*MVN(*t*) are baseline-subtracted forms of mossy fiber and MVN neuron firing. We assume that the timescale of the average indicated by the brackets is slow compared to variation in the input but faster than the timescale of plasticity of *v* (as for *τ*_*f*_ in the main text). Both before training and after consolidation, when *w* → 0, so that PC(*t*) → PC_0_,

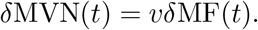

Then, the learning rule is at steady state when

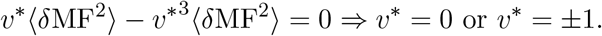

This is because Oja’s rule is designed to maintain the Euclidean length of the weight vector fixed at 1, and here there is only one input source with a plastic weight. Even in the case where there are multiple mossy fiber input types with plastic weights, Oja’s rule will drive the weight vector so that it is parallel (or antiparallel) to the eigenvector of the covariance matrix of the inputs that has largest eigenvalue (Oja 1982). In our model, in which we do not expect the signals carried by the mossy fiber inputs to themselves change persistently as a result of training, only one stable weight configuration can be maintained. Thus, Oja’s rule does not readily support consolidation in our model.

We next examine the Bienenstock-Cooper-Monroe (BCM) rule (Bienenstock *et al*. 1982), one of the most well-known firing rate-target homeostasis rules. The BCM rule consists of a Hebbian term multiplied by a term that changes the sign of plasticity depending on whether postsynaptic activity is greater than or less than a sliding threshold. For our circuit, this takes the form

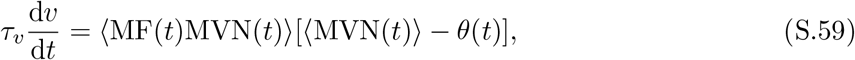

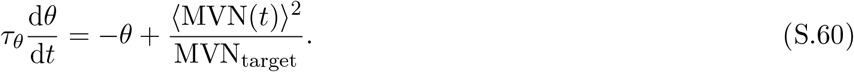

The angle brackets again represent time averages taken over a timescale longer than the variation in the input, but much shorter than the timescale of plasticity. The system has a fixed point whenever *θ* = ⟨MVN⟩ = MVN_target_.

There is one fixed point, with steady state weight value

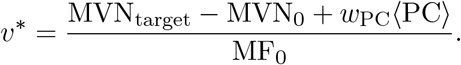

The value of *v*^∗^ is determined by parameters of the system that do not change as a result of learning (MF_0_, MVN_target_ and MVN_0_), as well as a variable (⟨PC⟩) that returns to its pre-training baseline value during the post-training period. Therefore, the system with one plastic synapse cannot persistently hold all possible values of *v* that may be reached after training.

For multiple plastic inputs to the (linear) MVN neuron, the BCM rule is less restrictive than Oja’s rule: there is a hyperplane—i.e., a continuum—in synaptic weight space of stable weight configurations for which the homeostatic condition of postsynaptic firing rate reaching a target is achieved. This opens the possibility that a more complex model of the circuit could allow consolidation of an analog memory using the BCM rule. However, such a rule would not allow for the weight of a single input type to change independently of the others, unlike the heterosynaptic rule.

## S5 Analysis of model dynamics with fixed-strength internal feedback loop

In this section, we analyze the dynamics of the model with a fixed-strength internal feedback loop (Materials and Methods, *Circuit model with internal feedback loop*). We first show the conditions under which stable post-training consolidation occurs. We then find conditions under which the circuit can also stably learn a target input-to-output gain during training.

### S5.1 Conditions for stable post-training consolidation

For stable consolidation, we need the late-learning site to stably hold any value of *v*. Following the same line of reasoning leading to Eq. (S.3), we can rewrite the heterosynaptic plasticity rule, Eq. (28), in terms of the weights as

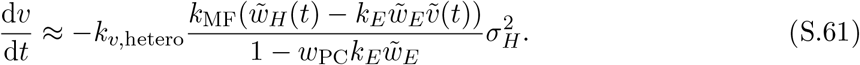

Note that, unlike the feedforward model, there must be variability in the post-training input, 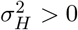, for consolidation to occur. From Eq. (S.61), *v* is at steady state as long as the feedforward weight at the early-learning weight is

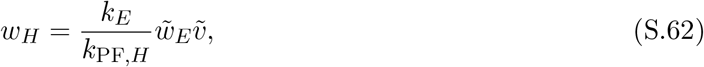

which defines a line in *w*_*H*_-*v* space (Fig. 6*E,J*). Values of *w*_*H*_ and *v* that lie along this line cause the early-learning area to not respond to sensory input to the circuit, i.e., the reset condition is satisfied. If *w*_*E*_ *>* 0, this can no longer be achieved in general by a passive decay at *w*_*H*_ to zero, as *w*_*H*_ needs to go to a positive steady state value since *v* is also positive. Below we evaluate the conditions under which this occurs for the two post-training reset mechanisms.

#### Inhibitory plasticity-driven resetting

In the inhibitory plasticity-driven reset mechanism, the weight of feedforward inhibition onto the early-learning area is governed by a Hebbian-like covariance rule, Eq. (29):

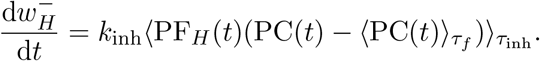

During the post-training period, the correlation between climbing fibers and the feedforward parallel fibers is zero, *c*_*H*_(*t*) ≡ 0, so simplifying as for Eq. (S.61), and combining with the learning rule for the excitatory weight, Eq. (27), we can write the equation for the overall feedforward weight at the early-learning site as

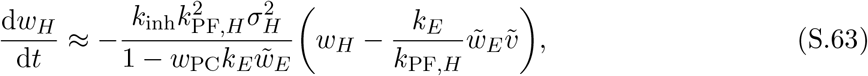

which has a fixed point that lies along the line in Eq. (S.62), satisfying the reset condition, as long as

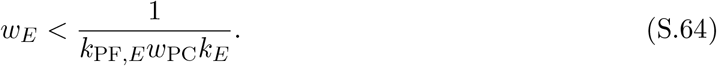

To determine whether the fixed point is attractive, it suffices to show that the slope of the trajectory approaching the fixed point is smaller than the slope of the line in Eq. (S.62). Otherwise, trajectories will move continuously away from the line of fixed points. From Eq. (S.63) and Eq. (S.61), trajectories have slope

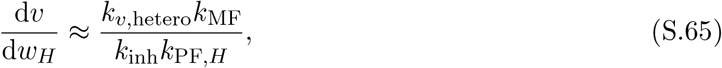

whereas from Eq. (S.62) the slope of the line of fixed points is 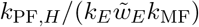. Thus, the slope of trajectories is smaller if

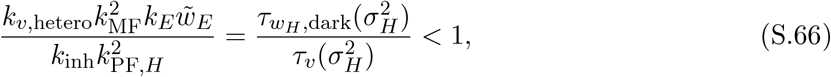

where the time constants 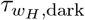 and *τ*_*v*_ are defined from Eq. (S.63) and Eq. (S.61) respectively as functions of the input variability 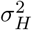,

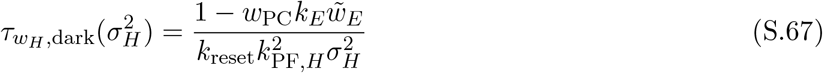

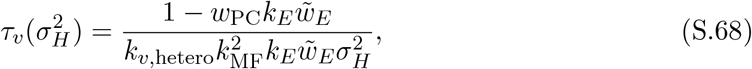

and where, to highlight the similarity of these expressions to the analogous quantities for the other reset mechanism below, we have defined

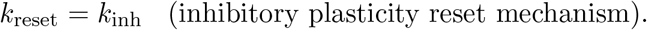

In other words, the line in Eq. (S.62) is an attractor only if plasticity is faster at the early-learning site than at the late-learning site.

The inhibitory plasticity here drives *w*_*H*_ so that the activity of the early-learning area does not modulate in response to sensory input. This result is broadly consistent with previous work showing that Hebbian plasticity of inhibitory synapses tends toward restoring excitatory-inhibitory balance (Vogels *et al*. 2011), except here this balance is only in terms of the fluctuating components of the inputs and not the spontaneous components, because of the sliding threshold term in Eq. (29). Note that, if 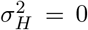, early-learning activity will not be reset, but there will also be no consolidation at *v*.

#### Inhibition of instructive signals-driven resetting

In the inhibition of instructive signals-driven reset mechanism, the activity of the pathway carrying instructive signals is inhibited by an output pathway from the late-learning area (Fig. 6*F* ; Eq. (30)):

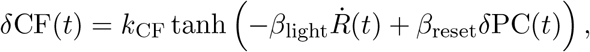

where 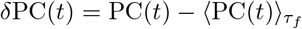. When information about errors is not present, 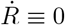 ≡ 0, so that for small errors, the parallel fiber-climbing fiber covariance *c*_*H*_(*t*) is in the linear regime, and the learning rule for *w*_*H*_ becomes

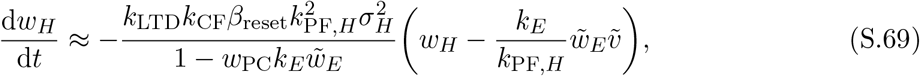

which has the same fixed points as Eq. (S.63) that satisfy the reset condition if Eq. (S.64) holds. Similarly, as in Eq. (S.66), the fixed point for a given trajectory is attractive if 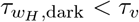 *< τ*_*v*_, where 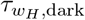 and *τ*_*v*_ are defined as in Eq. (S.67) and Eq. (S.68) but where we now define

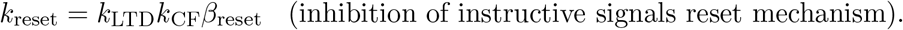

For large errors, the magnitude of *δ*CF will be saturated, and d*w*_*H*_*/*d*t* will reach a maximum value, so the slope of trajectories will be

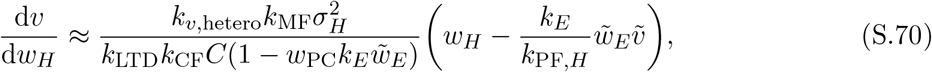

where *C* is defined as in Eq. (S.5). That is, the trajectories become increasingly vertical the further they are from the line of fixed points. For trajectories to converge to a stable fixed point, they must have slopes shallower than the line of fixed points. This occurs when

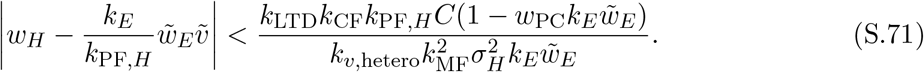

This effectively places a limit on how large a gain change the circuit can learn during the training period in order for that gain change to be stably consolidated (see §S5.2, *Inhibition of instructive signals-driven resetting*).

### S5.2 Dynamics during training

Here we analyze the dynamics of the synaptic weights in the circuit during training for each mechanism, and show that under similar conditions as post-training, both mechanisms are also stable during training and result in the circuit correctly learning the target input-to-output computation (Fig. S3).

#### Local stability of the fixed point during training

We first show that both mechanisms have the same locally stable fixed point, corresponding to the circuit achieving the target gain. For both mechanisms, the learning rule for the late-learning weight *v* is identical, Eq. (S.61), and the only difference comes from the effect on the early-learning feedforward weight *w*_*H*_. We assume that the weight of the feedback pathway is fixed and that the total feedback strength is less than 1, as in Eq. (S.64). We also assume that 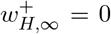 aand that during training the variability of the head input is 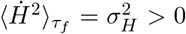. In our analysis, we make use of the fact that, from Eq. (23), the error in gain can be expanded as

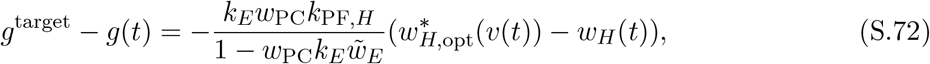

and for convenience we use variables with tildes as defined in Eq. (24)–Eq. (26), and where

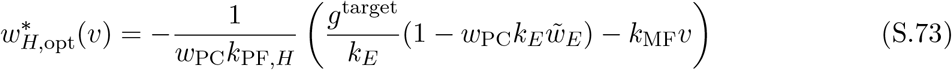

is the value of *w*_*H*_ that would minimize the error in gain for a fixed choice of *v*. Similarly, we define the value of *w*_*H*_ that would reset early-learning activity for a fixed choice of *v* as

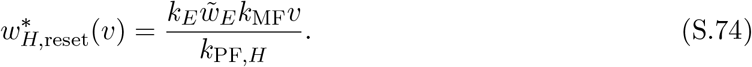

For the inhibitory plasticity mechanism (Fig. 6*A*), changes in the excitatory weight 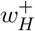 are governed by instructive climbing fiber input driven by the retinal slip signal, Eq. (12), according to the learning rule in Eq. (27), whereas the inhibitory weight 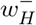 tends to drive the activity of the early-learning area back to its long-term average, as in Eq. (29). More precisely, for a target gain value *g*^target^, plasticity at 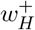 is driven by the parallel fiber-climbing fiber covariance

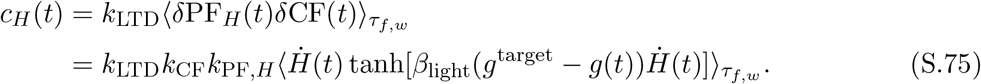

We can combine the learning rules for 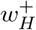 and the inhibitory weight 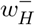 to write an expression for the change in the net weight 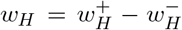, and simplify using the same logic as for the post-training period (see §S1). This yields

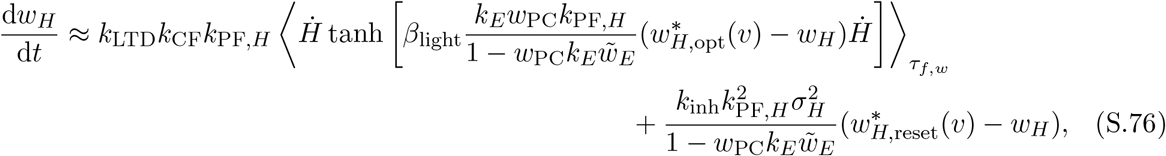

using the simplifications in Eqs. (S.72)–(S.74).

For the inhibition of instructive signals mechanism (Fig. 6*F*), the climbing fiber input is both driven by retinal slip error and inhibited by a pathway carrying Purkinje cell output, Eq. (30), leading to a parallel fiber-climbing fiber covariance

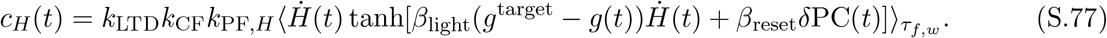

This yields a simplified learning rule for *w*_*H*_:

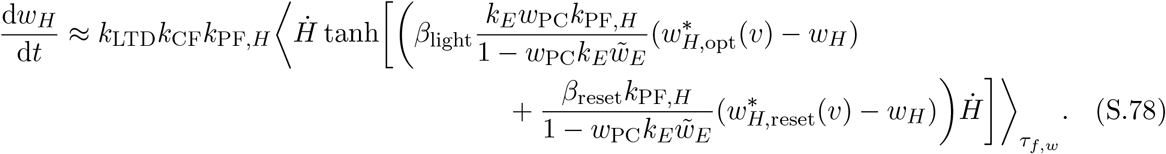

Both mechanisms have the same fixed point,

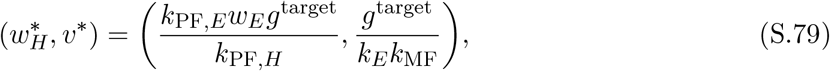

at which point the gain is equal to *g*^target^ and 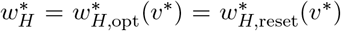. Furthermore, the fixed point lies along the post-training line attractor, Eq. (S.62). Close to the fixed point, the dynamics for both mechanisms are linear, as the hyperbolic tangent term is in an approximately linear regime,

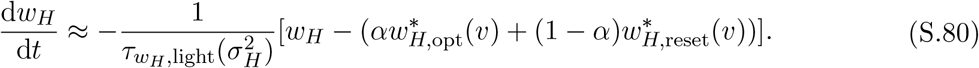

where we defined

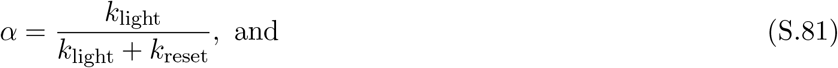

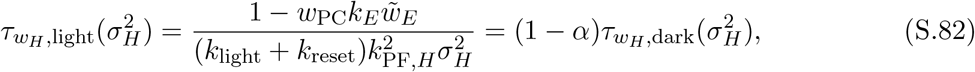

with

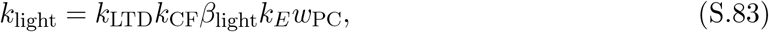

and

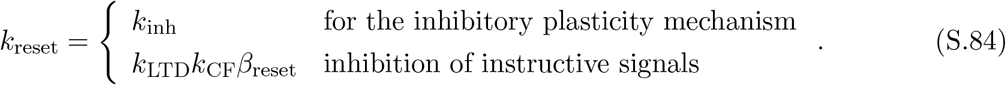

From Eq. (S.61) and Eq. (S.80), the Jacobian is

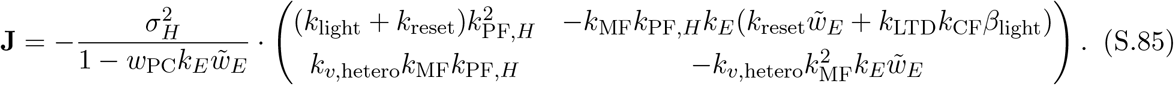

The eigenvalues of the Jacobian are

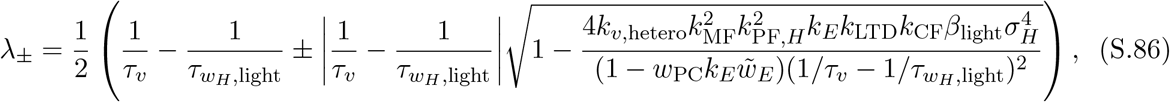

where for conciseness, we omitted that 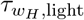 and *τ*_*v*_ are functions of 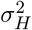, with the latter defined as in Eq. (S.68). Since by definition all parameter values are positive, during training 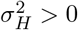, and we assumed that Eq. (S.64) holds, both eigenvalues have negative real part and the fixed point is stable if

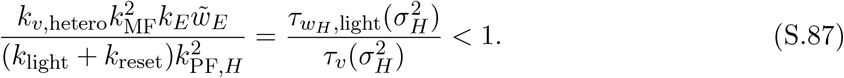

Note that if the post-training stability condition, Eq. (S.66), is met, then this stability condition is also automatically met, because by definition 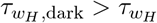 for all 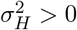 (see Eq. (S.82)).

Outside of the region near the fixed point, the dynamics of the weights differ between the two mechanisms. In the following sections, we evaluate the stability of the dynamics in this region for each mechanism.

#### Inhibitory plasticity-driven resetting

For the inhibitory plasticity mechanism, we can understand the dynamics by first looking at *w*_*H*_, as defined by the learning rule in Eq. (S.76). During training to increase the gain, *w*_*H*_ will both be driven by the climbing fiber input away from its initial value along the post-training attractor, towards the line of points 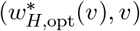 (along which *g* = *g*^target^), and by the inhibitory plasticity back towards the attractor (the *v*-nullcline). Thus, the nullcline for *w*_*H*_ lies in between these two lines. As training continues, the weight trajectory will cross the *w*_*H*_-nullcline, entering a region in which both *v* and *w*_*H*_ are increasing. If the slope of the flow field along the 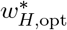 line is shallower than the slope of the line itself, then the trajectory will stay bounded between the 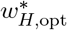 line and the *w*_*H*_-nullcline, reaching the fixed point without any overshoot in the circuit gain (Fig. S3*A,B*). Otherwise, the trajectory will cross the 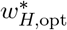 line, causing an overshoot in the gain. In this case, the trajectory will still tend toward the fixed point as long as the post-training stability condition, Eq. (S.66), holds.

First, we determine the conditions under which the trajectory will reach the fixed point without any overshoot in gain. The slope of the flow field along the 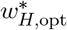 line is

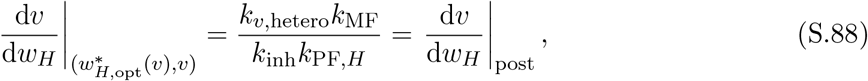

i.e., it is equal to the slope of trajectories during the post-training period (see Eq. (S.65)). The slope of the 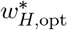 line is

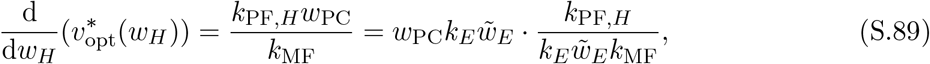

where we rearrange Eq. (S.73) to define 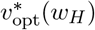 as the value of *v* that satisfies 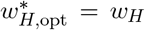. Note that the second term in the rightmost expression above is the slope of the *v*-nullcline (i.e., the post-training attractor). Therefore, following the reasoning leading to the post-training stability condition in Eq. (S.66), the slope of the flow field along the 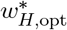 line will be shallower than the slope of the line itself if

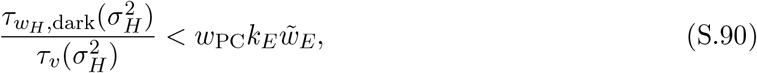

which can also be written as

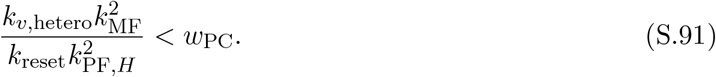

Note that, since 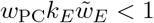 (Eq. (S.64)), this is a stricter condition than Eq. (S.66).

If the condition in Eq. (S.90) does not hold, then the trajectory will cross the 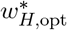 line. In this region, from Eq. (S.61) and Eq. (S.72)–Eq. (S.76), the instantaneous slope of the trajectory is given by

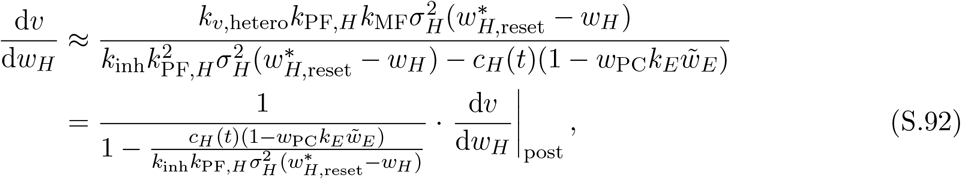

where *c*_*H*_(*t*) is as defined in Eq. (S.75). Since *c*_*H*_ has a saturating nonlinearity, −*Ck*_LTD_*k*_CF_*k*_PF,*H*_ *< c*_*H*_ ≤ 0 in the region 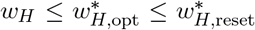, with *C* as defined by Eq. (S.5). Then, the slope of trajectories is bounded,

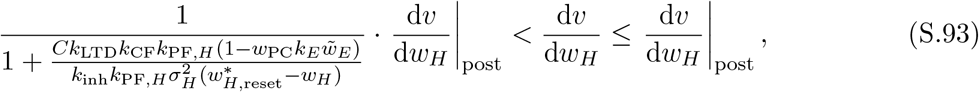

where the right-hand equality holds along the 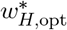 line, as we saw above. Therefore, as long as Eq. (S.66) holds (i.e., the slope of post-training trajectories is shallower than the slope of the *v*-nullcline), trajectories during training will also have slope shallower than the *v*-nullcline and tend toward the fixed point.

#### Inhibition of instructive signals-driven resetting

For the inhibition of instructive signals mechanism, we saw above (§S5.1, *Inhibition of instructive signals-driven resetting*) that post-training dynamics could become unstable if the change in *w*_*H*_ during the training period was too large, limiting the maximum change in gain that the circuit could stably consolidate. This was defined by the slope of trajectories in the region of weight space where the climbing fiber response was saturated, Eq. (S.70), from which we found a stable region of weight space, Eq. (S.71). Assuming that before training the weights start at a fixed point and that during training the learned change in gain initially only comes from changes in *w*_*H*_, the largest gain change that could be learned is

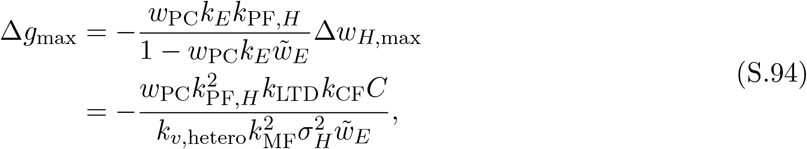

where Δ*w*_*H*,max_ has value equal to the right hand side of Eq. (S.71). With this largest gain change in mind, below we examine the weight dynamics during training.

From Eq. (S.78) we can see that for a fixed value of *v*, d*w*_*H*_*/*d*t* = 0 when

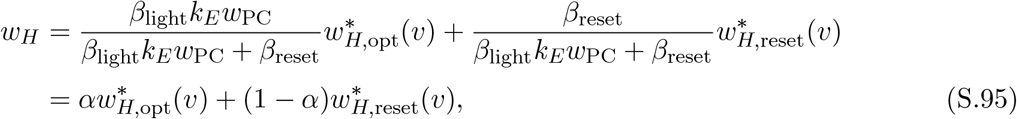

where *α* is defined as in Eq. (S.81). That is, the *w*_*H*_-nullcline is linear and lies between the line 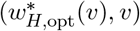 (along which *g* = *g*^target^) and the post-training attractor (*v*-nullcline). Thus, during training to increase the gain, initially *w*_*H*_ will decrease from a point on the post-training attractor toward the *w*_*H*_-nullcline, during which time *v* will start to grow. The trajectory will then cross the *w*_*H*_-nullcline. As long as the trajectory does so within the region defined by Eq. (S.71), the trajectory will tend toward the post-training attractor and eventually the fixed point (Fig. S3*D*).

However, it is possible that trajectories may cross the 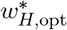 line, which would correspond to the gain of the circuit overshooting the target. We can try to ensure that trajectories do not overshoot by keeping them within the region bounded by 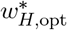 and the *w*_*H*_-nullcline. If the 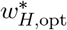 line is near the *w*_*H*_-nullcline and therefore within the linear regime of the climbing fiber response *δ*CF, the slope of trajectories along the line is approximately as defined in Eq. (S.89). As for the inhibitory plasticity reset mechanism, trajectories will therefore stay approximately within the region between the 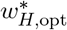 line and the *w*_*H*_-nullcline if the condition in Eq. (S.90) is met. If the 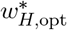 line is far from the nullcline, and the climbing fiber response is saturated, then the slope of the flow field along the 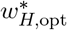 line is

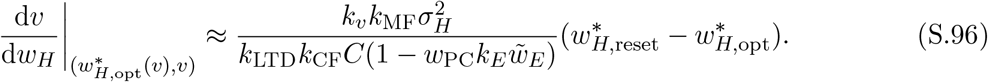

Then, trajectories will stay bounded by the 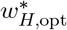 line and there will be no overshoot in gain if

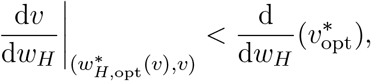

where 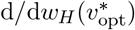 is the slope of the 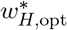 line defined in Eq. (S.89). This directly gives that, for a given value of *v*, the distance between the 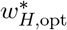 line and the *v*-nullcline 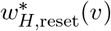(*v*) can be no larger than

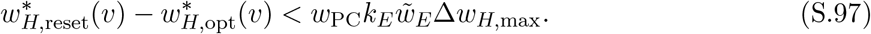

### S5.3 Effect of post-training reset on weight dynamics

With either the inhibitory plasticity or homeostatic climbing fiber mechanism, two important properties arise in the model’s dynamics that are also observed experimentally. First, the early-learning weight *w*_*H*_ tends toward a saturation point during training that is not simply the value that would minimize errors. In the feedforward model, this property was modeled explicitly by including a decay term in the learning rule Eq. (13) (following previous work), which we do not include in the model with internal feedback (i.e., in Eq. (27)). Second, during training, the approach of the early-learning weight to the saturation point occurs over a faster timescale than the subsequent post-training reset during consolidation (i.e., Eq. (S.82); for the inhibition of instructive signals, this is true in the linear regime of the CF response). This was also modeled explicitly in the feed-forward model with two different values of *τ*_*w*_ during the training and post-training periods, but in the model with internal feedback results from the fact that learning and resetting are being driven by two different mechanisms. Note that we could also include either of the reset mechanisms in the feedforward model (as it is the case of *w*_*E*_ = 0), and the resulting dynamics would also have these properties without having to explicitly model them.

## S6 Circuit model with plastic internal feedback

In this section, we extend the feedback model discussed in the previous section (§S5) to the case in which the weight of the internal feedback pathway to the early-learning area is also plastic. Plasticity in the excitatory feedforward and feedback weights, 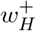 and 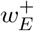 respectively, is described by Eq. (27). The plasticity at the late-learning site *v* is described, as before, by Eq. (S.61). Similar to the circuits with fixed strength feedback, we can show how changes at the early-learning site can be successfully consolidated post-training if the circuit either has plasticity at 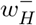 (and/or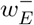), or through inhibition of the climbing fiber instructive signal pathway, both of which reset early-learning activity so that the output of the early-learning area does not modulate in response to sensory input.

Below, we describe in further detail the implementation of each of these reset mechanisms in the circuit. Then, we show that the linearized dynamics of the circuit with either reset mechanism are the same, and determine conditions under which learning and consolidation occur stably. In general, learning of a target input-output gain and post-training consolidation will be locally stable around its fixed point as long as the weight of the feedback pathway *w*_*E*_ stays within the region *w*_*E*_ *<* 1*/*(*w*_PC_*k*_*E*_*k*_PF,*E*_), so that the strength of the feedback around the loop from early-to late-learning area and back to the early-learning area is less than 1, and if the rate of plasticity at the early-learning sites is fast compared to the rate of plasticity at the late-learning site.

### S6.1 Model formulation

We implemented the two circuit reset mechanisms by extending the model with non-plastic feedback (see Materials and Methods, *Circuit model with internal feedback loop*) as follows.

#### Inhibitory plasticity-driven resetting

For the inhibitory plasticity mechanism we model plasticity in 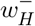 as governed by Eq. (29) and in 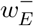 by

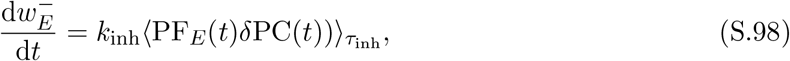

Where 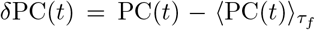. During training with a target gain of *g* ^target^, plasticity at the excitatory weights to the early-learning area is driven by a climbing fiber signal carrying only retinal slip, Eq. (12). Plasticity at the feedforward excitatory weight 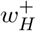 is driven by (negative of) the feedforward parallel fiber-climbing fiber covariance *c*_*H*_(*t*), Eq. (S.75), leading to the simplified learning rule for the net weight *w*_*H*_ in Eq. (S.76), where now 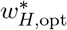 and 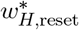 are functions of both *w*_*E*_ and *v*. Similarly, plasticity at 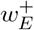 is driven by (negative of) the feedback covariance

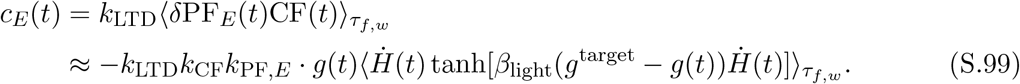

As for the feedforward input, we note that

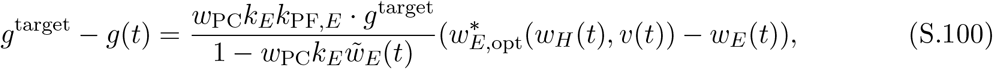

where we define

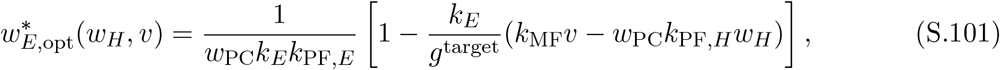

i.e., the value of *w*_*E*_ that would minimize errors for a fixed choice of *w*_*H*_ and *v*. Further defining

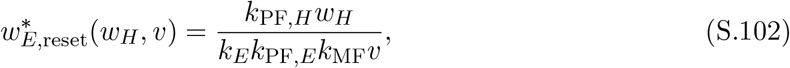

we can write a combined learning rule for the net feedback weight *w*_*E*_,

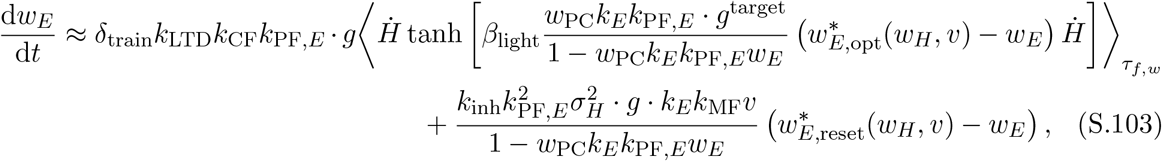

where *δ*_train_ = 1 during training. Post-training, *δ*_train_ = 0 since 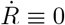, which implies that *c*_*H*_(*t*) = *c*_*E*_(*t*) ≡ 0. Similarly, post-training plasticity at *w*_*H*_ is governed by Eq. (S.63).

#### Inhibition of instructive signals-driven resetting

For the inhibition of instructive signals mechanism, we assume the inhibitory weight is fixed and that plasticity at both 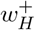 and 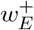 is governed by the climbing fiber response defined in Eq. (30) (Fig. S2*A*). More specifically, plasticity at 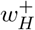 is driven by *c*_*H*_(*t*) as defined in Eq. (S.77), and we can similarly write the feedback parallel fiber-climbing fiber covariance that drives plasticity at 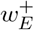 as

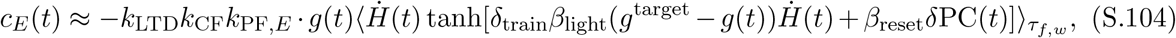

where *g*^target^ is the target gain of the circuit during training, and *δ*_train_ = 1 during training and 0 post-training. From this, we can write that plasticity of the net weight *w*_*E*_ is governed by

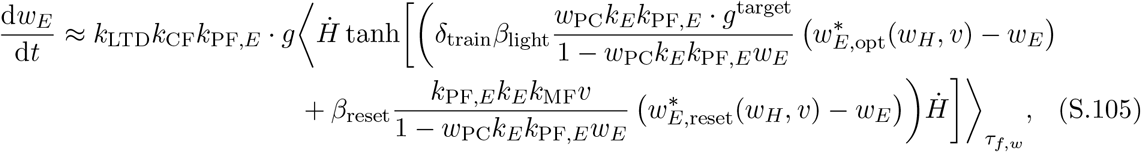

where 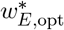 and 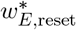 are defined by Eq. (S.101) and Eq. (S.102) above.

In Figure S2, we simulated the time evolution of the weights in the circuit implementing this latter mechanism for a 0.5 h period of training to increase the gain to *g*^target^ = 2 followed by a 23.5 h post-training period. The model parameters were the same as for the model without plasticity of the feedback pathway (Tables S1–S3; see Materials and Methods, *Simulation of oculomotor learning*).

### S6.2 Stability of shared linearized dynamics

Here we extend the analyses of the model without plastic feedback during the training (§S5.2) and post-training (§S5.1) periods to the case of plastic feedback.

#### Training

We consider the system of differential equations for the three synaptic weights *w*_*H*_, *w*_*E*_ and *v* from Eq. (S.61) and either Eq. (S.76) and Eq. (S.103) for the inhibitory plasticity reset mechanism, or Eq. (S.78) and Eq. (S.105) for the inhibition of instructive signals reset mechanism. For either mechanism, there is a line of steady states (parameterized by *s*),

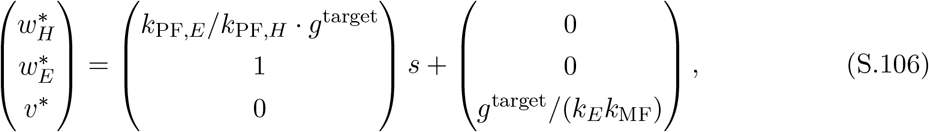

along which 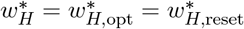 and 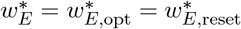. This corresponds to the intersection of the surface in weight space that leads the circuit to produce the target gain *g*^target^ with the surface defined by Eq. (S.62).

To analyze the stability of the fixed points defined by this line, we calculate the eigenvalues of the Jacobian of the system of differential equations. We note that the Jacobian has the same form for both reset mechanisms after substituting the relevant expressions for *k*_reset_, as defined in Eq. (S.84), into the differential equations. Leaving Eq. (S.106) parameterized by 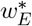 (i.e., taking an arbitrary value of *s*), we calculate the characteristic polynomial of the Jacobian, which has three roots. One of the roots is *λ*_1_ = 0 and corresponds to the direction along the line of steady states. Dividing this root out, we are left with *λ*^2^ + *aλ* + *b* = 0 for determining the remaining eigenvalues *λ*_2_ and *λ*_3_, where

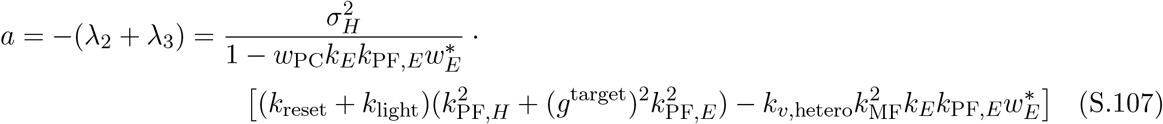

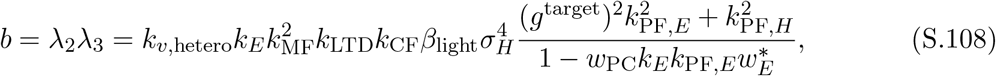

using *k*_light_ and *k*_reset_ as defined in Eqs. (S.83) and (S.84). For the eigenvalues to have negative real parts, we must have *a >* 0 and *b >* 0. For *w*_*E*_ *<* 1*/*(*w*_PC_*k*_*E*_*k*_PF,*E*_), since all parameters are positive and 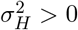 during training, *b >* 0 is satisfied. *a >* 0 is satisfied if

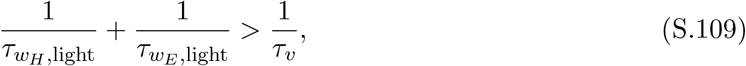

where 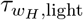 and *τ*_*v*_ are defined as in Eqs. (S.82) and (S.68) and depend on the value of 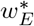 parameterizes the fixed point as well as on 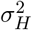, and where

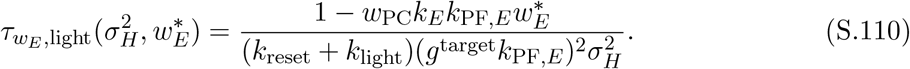

That is, for stable learning, the sum of the instantaneous effective rates of plasticity of the early-learning sites *w*_*H*_ and *w*_*E*_ should be faster than the instantaneous effective rate of plasticity at the late-learning site *v*, extending the result from the fixed-strength feedback case above.

Assuming that 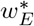 is upper-bounded by 1*/*(*w*_PC_*k*_*E*_*k*_PF,*E*_), and assuming that the circuit is learning only positive gains so that *g*^target^ *>* 0, we can satisfy Eq. (S.109) for the whole range of 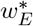 and potential gains with a stricter condition

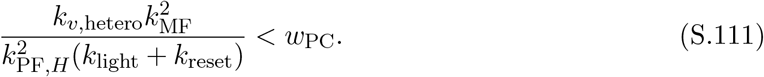

#### Post-training

In the absence of information about errors, we take *δ*_train_ = 0 in Eqs. (S.103) and (S.105). Then, the steady states of the dynamics lie along the surface represented by Eq. (S.62).

Calculating the eigenvalues of the Jacobian at a fixed point 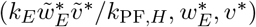, parameterized by 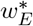 and *v*^∗^, we find that there are two zero eigenvalues, corresponding to the surface in Eq. (S.62), and a third eigenvalue

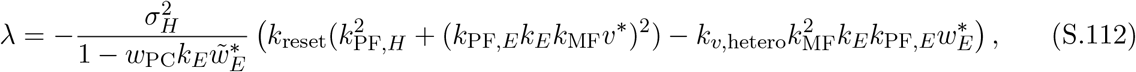

which is negative if 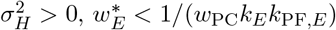 and

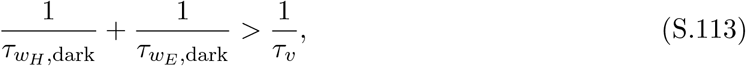

where 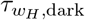 and *τ*_*v*_ are defined as in Eq. (S.67) and Eq. (S.68) and depend on the value of 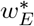 that parameterizes the fixed point as well as on 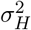, and where

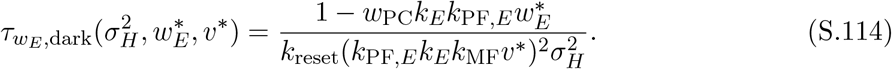

For 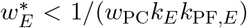, and assuming *v*^∗^ *>* 0, we can write a single stricter condition that is equivalent to Eq. (S.91) to satisfy Eq. (S.113). If this condition is met, then the surface of fixed points Eq. (S.62) will be locally attractive during the post-training period, and since Eq. (S.111) will be automatically satisfied, the dynamics during training will be locally stable around the fixed point as well.

**Figure S1:**
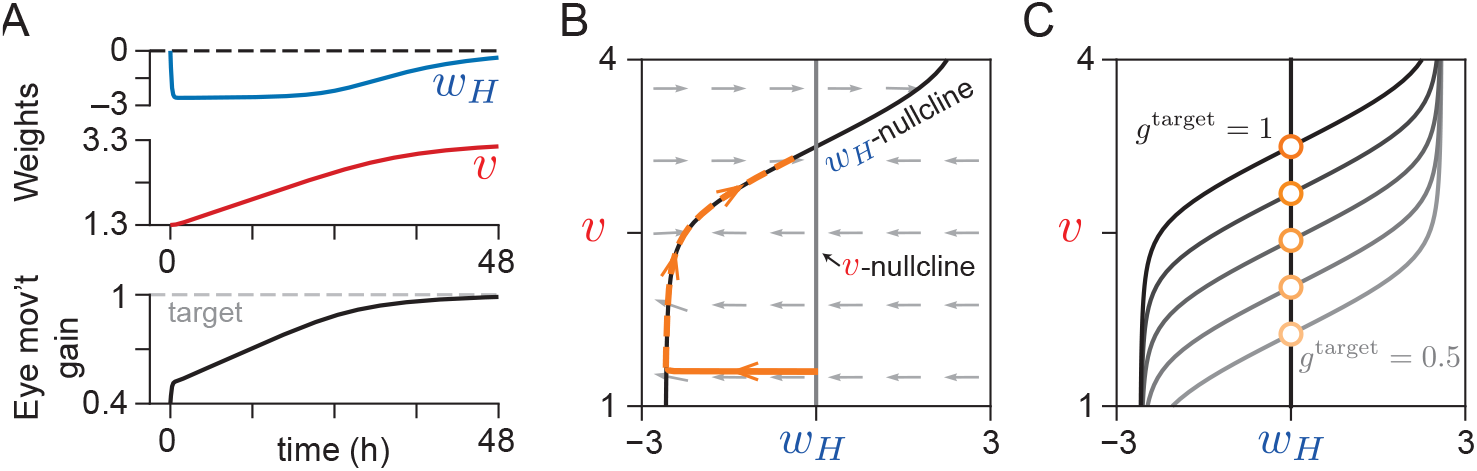
Synaptic weight dynamics of the feedforward-architecture model during training. (*A*) Dynamics of weights (*top*) and the gain of the circuit (*bottom*) during 48 h of simulated training to increase the sensory input-to-motor output gain of the circuit. (*B*) Trajectory (orange curve) shows the evolution of synaptic weights in *A*, which approaches the stable fixed point where the *w*_*H*_ - and *v*-nullclines (light and dark grey lines) cross. For clarity, the orange trajectory is shown as dashed where it overlaps the *w*_*H*_ -nullcline. (*C*) The nullclines (grey to black lines) and fixed points (orange circles) of the system for different choices of desired input-to-output gain value (shown by lightness). Any target gain value can be learned by the system, corresponding to a fixed point lying along the line *w*_*H*_ = 0. These points are thus also stable when information about behavioral errors is not present.

**Figure S2:**
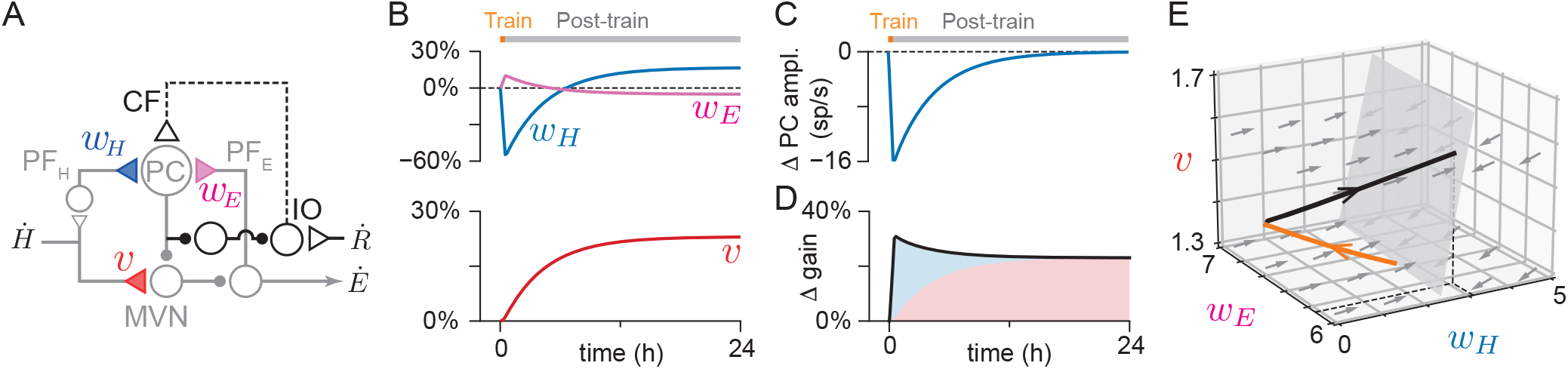
Systems consolidation in a circuit with plastic internal feedback loop. (*A*) Architecture of circuit with three sites of plasticity, at the feedforward weight onto the early-learning area, *w*_*H*_ (blue), at the weight of the internal feedback pathway, *w*_*E*_ (pink), and at the direct pathway onto the late-learning area, *v* (red). As in Fig. 6*F*, the post-training reset was achieved by inhibition of the pathway carrying instructive signals to the early-learning area. (*B*) Change in weights during 0.5 h of simulated training to increase the gain (orange block) followed by 23.5 h post-training with no feedback about behavioral errors (grey block). (*C*) Early-learning area output (amplitude of Purkinje cell activity relative to moving baseline), which drives consolidation at the late-learning site. (*D*) Change in the gain of the eye movement response (black line). Blue and red shaded areas show the contributions of the early- and late-learning areas to the circuit transformation. (*E*) Trajectory of synaptic weights during the training (orange) and post-training (black) periods. Dashed black lines show the projection of the steady state reached at the end of training onto the *w*_*H*_ -*w*_*E*_ plane. Grey arrows show the approximate instantaneous direction in which a weight configuration at a given point in synaptic weight space will evolve during the post-training period, determined analytically. Trajectories tend toward a 2-D surface of marginally stable points (solid grey surface).

**Figure S3:**
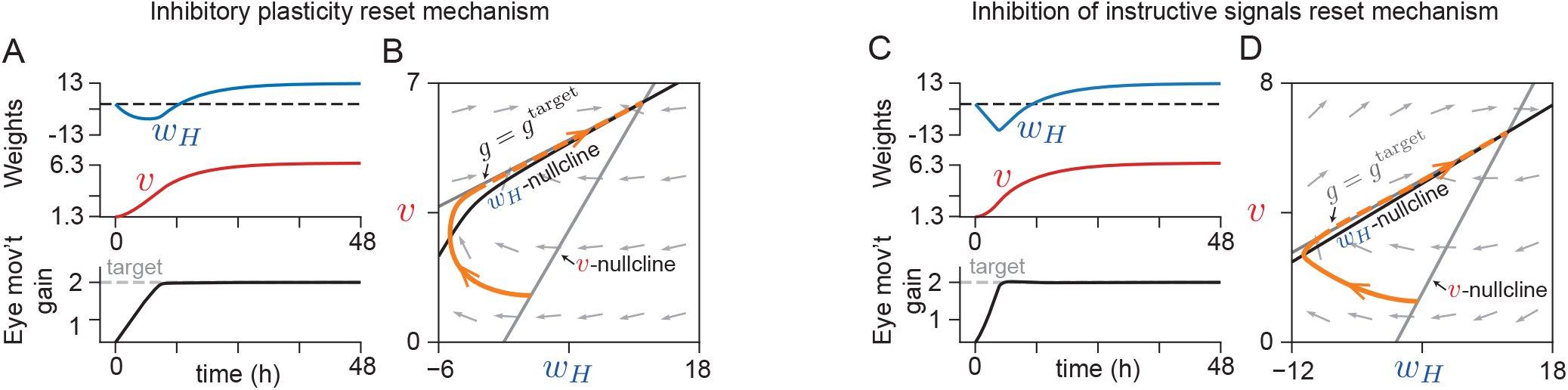
Synaptic weight dynamics during training of the model with feedback architecture. (*A*) Dynamics of weights (*top*) and the gain of the circuit (*bottom*) during 48 h of simulated training to increase the sensory input-to-motor output gain of the circuit, using the inhibitory plasticity reset mechanism. (*B*) Trajectory (orange curve) shows the evolution of synaptic weights in *A*. All trajectories approach a stable fixed point where the *w*_*H*_ - and *v*-nullclines cross. The second grey line that the trajectory follows corresponds to the constant gain line for the target gain. (*C,D*) Same as *A,B*, but for the inhibition of instructive signals reset mechanism. In panels *B* and *D*, for clarity, the orange trajectory is shown as dashed where it overlaps the *w*_*H*_ -nullcline.

**Table S1:** Values and descriptions for parameters of feedforward circuit model. Sources for values taken from the literature are provided in the right-hand column. Parameters that were calculated based on published values are indicated as “based on” a source. Table S1 continued.

| Parameter | Value | Description | Source |
| --- | --- | --- | --- |
| $MF_0$ | 55 sp/s | Mossy fiber (MF) spontaneous baseline rate | Lasker *et al.* (2008) |
| $k_{MF}$ | 0.14 (sp/s)/(deg/s) | MF sensitivity to vestibular input | |
| $PF_0$ | 14 sp/s | Parallel fiber (PF) spontaneous baseline rate | Based on Arenz *et al.* (2008) |
| $k_{PF}$ | 0.42 (sp/s)/(deg/s) | PF sensitivity to vestibular input | |
| $PC_0$ | 50 sp/s | Purkinje cell spontaneous baseline rate | Kato *et al.* (2015) |
| $MVN_0$ | −12 sp/s | Correction factor to make medial vestibular nucleus (MVN) spontaneous baseline rate = 57 sp/s | Beraneck & Cullen (2007) |
| $CF_0$ | 1 sp/s | Climbing fiber (CF) spontaneous baseline rate | E.g., Goossens *et al.* (2004) and Maruta *et al.* (2007) |
| $k_{CF}$ | 1 sp/s | Maximum amplitude of CF response around baseline | |
| $\beta_{light}$ | 1 s/deg | Scale factor for saturation of CF response | |
| $k_E$ | 2.2 (deg/s)/(sp/s) | Eye velocity sensitivity to MVN firing | Based on Beraneck & Cullen (2007) |
| $k_{LTP}$ | 1.005 s/sp | Contribution of parallel fiber firing to plasticity at early-learning site $w_H$ | Based on amount of learning in Kimpo *et al.* (2014) and Boyden & Raymond (2003) |
| $k_{LTD}$ | 0.648 (s/sp)<sup>2</sup> | Contribution of parallel fiber-climbing fiber coincidence to plasticity at $w_H$ | |
| $\tau_{w,train}$ | 0.15 h | Time constant of plasticity at $w_H$ during training | Boyden & Raymond (2003) |
| $\tau_{w,post}$ | 5 h | Time constant of plasticity at $w_H$ after training | Based on Cooke *et al.* (2004) and Jang *et al.* (2020) |
| $k_{v,hetero}$ | $2.75 \times 10^{-5}$ (s/sp)<sup>2</sup>/h | Plasticity rate at late-learning site $v$ for heterosynaptic rule | |
| $k_{v,Hebb}$ | $8 \times 10^{-3}$ (s/sp)<sup>2</sup>/h | Plasticity rate at $v$ for Hebbian rule | |

Table S1 continued.
| Parameter | Value | Description | Source |
| --- | --- | --- | --- |
| $\tau_f$ | 1 min | Timescale of fast average used to calculate eye movement output | |
| $\tau_{f,w}$ | 1 min | Timescale of average used in plasticity rule at $w_H$ | |
| $\tau_{f,v}$ | 1 min (Fig. 2)<br>0.7 h (Fig. 3) | Timescale of average used in plasticity rule at $v$ | |
| $\tau_s$ | 0.0395 h | Timescale of sliding average $\theta$ in Hebbian rule for $v$ | |
| $w_H^-$ | 5 | Nonplastic molecular layer interneuron to Purkinje cell weight | Consistent with weight change in Jang *et al.* (2020) |
| $w_{H,0}^+$ | 5 | Initial value of early-learning, PF-Purkinje cell weight $w_H$ | |
| $w_{PC}$ | 0.05 | Purkinje cell to MVN synaptic strength | Based on Payne *et al.* (2019) |
| $v_0$ | 1.3 | Initial value of late-learning MF-MVN weight $v$ | |

**Table S2:**
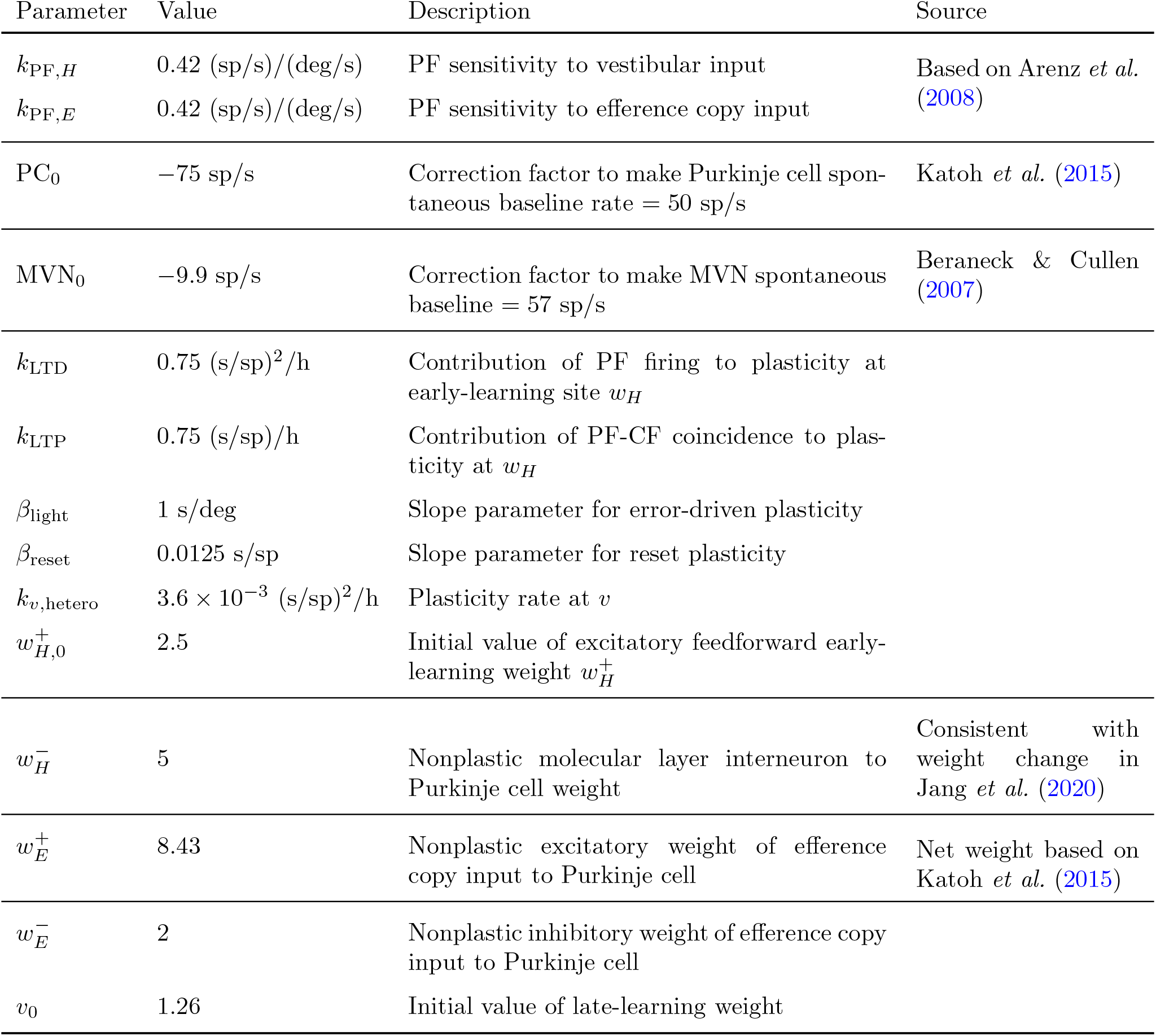
Values, descriptions and sources for parameters of circuit model with internal efference copy feedback, using the climbing fiber negative feedback reset mechanism. All other parameters same as in Table S1.

**Table S3:**
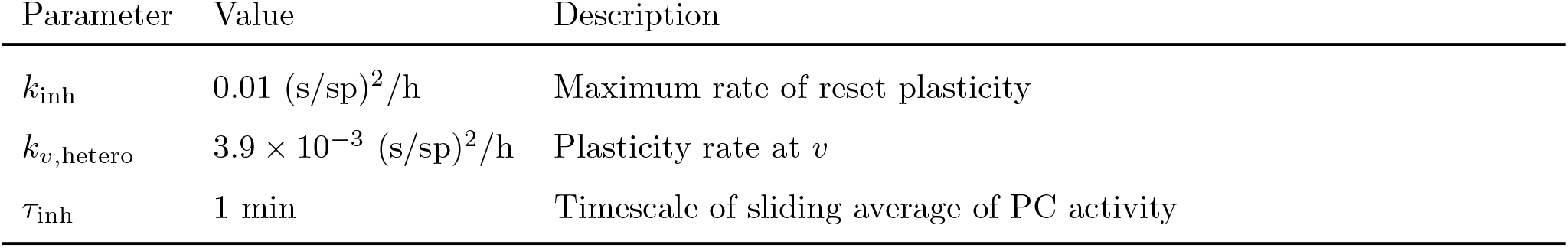
Values and descriptions of parameters of circuit model with internal efference copy feedback, using the inhibitory plasticity reset mechanism. All other parameters same as in Tables S1 and S2.

